# Type I interferon potentiates metabolic dysfunction, inflammation, and accelerated aging in mtDNA mutator mice

**DOI:** 10.1101/2020.09.22.308171

**Authors:** Yuanjiu Lei, Camila Guerra Martinez, Sylvia Torres-Odio, Samantha L. Bell, Christine E. Birdwell, Joshua D. Bryant, Carl W. Tong, Robert O. Watson, Laura Ciaccia West, A. Phillip West

**Affiliations:** Department of Microbial Pathogenesis and Immunology, College of Medicine, Texas A&M University Health Science Center, Bryan, Texas, USA; Department of Medical Physiology, College of Medicine, Texas A&M University Health Science Center, Bryan, Texas, USA

## Abstract

Mitochondrial dysfunction is a key driver of inflammatory responses in human disease. However, it remains unclear whether alterations in mitochondria-innate immune crosstalk contribute to the pathobiology of mitochondrial disorders and aging. Using the polymerase gamma (POLG) mutator model of mitochondrial DNA (mtDNA) instability, we report that aberrant activation of the type I interferon (IFN-I) innate immune axis potentiates immunometabolic dysfunction, reduces healthspan, and accelerates aging in mutator mice. Mechanistically, elevated IFN-I signaling suppresses activation of nuclear factor erythroid 2-related factor 2 (Nrf2), which increases oxidative stress, enhances pro-inflammatory cytokine responses, and accelerates metabolic dysfunction. Ablation of IFN-I signaling attenuates hyper-inflammatory phenotypes by restoring Nrf2 activity and reducing aerobic glycolysis, which combine to lessen cardiovascular and myeloid dysfunction in aged mutator mice. These findings further advance our knowledge of how mitochondrial dysfunction shapes innate immune responses and provide a framework for understanding mitochondria-driven immunopathology in POLG-related diseases and aging.

## Introduction

An expanding body of literature indicates that mitochondria are key regulators of the mammalian innate immune response, with both beneficial and deleterious consequences for the host (*1*). Mitochondria serve as antiviral signaling hubs and facilitate antibacterial immunity by generating reactive oxygen species (ROS), but they can also promote inflammation following cellular damage and stress (*2–5*). Recent work has demonstrated that mtDNA is a potent agonist of nucleic acid sensors of the innate immune system, including Toll-like receptor 9 (TLR9), NLR family, Pyrin domain containing 3 (NLRP3), and cGAS (*6, 7*). The cGAS-STING axis is now recognized as a major driver of IFN-I and inflammatory responses to nuclear and mitochondrial genome instability, and the aberrant release of mtDNA from damaged cells and tissues is increasingly linked to a growing list of human diseases (*6, 8–10*).

Mitochondrial diseases (MDs) are a group of clinically heterogeneous disorders caused by inherited mutations in genes that function in oxidative phosphorylation (OXPHOS) and mitochondrial metabolism (*11, 12*). In addition to exhibiting metabolic and energetic deficits, patients with MDs are more susceptible to opportunistic pathogens and also suffer elevated complications arising from these infections (*13–15*). Although B and T cell immunodeficiencies can contribute to recurrent infections in MDs (*16*), comparatively little is known about innate immune dysregulation in patients and/or murine models. Hyperactivation of the innate immune system is a key feature of sepsis, systemic inflammatory response syndrome (SIRS), and acute respiratory distress syndrome, all of which occur more frequently and lead to significant mortality in patients with MD (*14, 15*). Given the established and emerging links between mitochondria and innate immunity, persistent mitochondrial dysfunction in MD could basally activate or rewire the innate immune system. This could occur as loss of mitochondrial integrity and/or quality control liberates mitochondrial damage-associated molecular patterns (DAMPs), such as mtDNA, which engage the innate immune signaling and promote inflammatory responses that synergize with metabolic impairments to drive pathology. Accordingly, elevated inflammatory cytokines have been observed in patients with Alpers-Huttenlocher syndrome and mouse models of primary mitochondrial disorders, suggestive of heightened innate immune activation (*17, 18*).

The mitochondrial polymerase gamma (POLG) enzyme possesses DNA polymerase and 3’→5’ DNA exonuclease activities, and nearly 250 pathogenic mutations in *POLG* have been linked to diseases including primary MDs, parkinsonism, and cancer (*19–22*). Mutations in *POLG* represent the most prevalent single-gene cause of MD, and are implicated in a range of disorders including Alpers-Huttenlocher syndrome, ataxia neuropathy spectrum, and progressive external ophthalmoplegia (PEO), all of which are characterized by multiple organ pathology with varying degrees of nervous, muscular, digestive, and endocrine system involvement. In recent years, several mouse models of POLG-related disease have been reported, the most well studied of which is the POLG mutator mouse (*23, 24*). These knock-in mice contain D257A substitutions in the exonuclease domain, and animals homozygous for the mutant alleles exhibit disrupted exonuclease function and elevated mtDNA instability. POLG mutator mice present pathology that mirrors various aspects of human MDs, including cardiomyopathy, progressive anemia, and sensorineural hearing loss. These animals also display premature aging between 6 and 9 months, characterized by alopecia, osteoporosis, kyphosis, and decreased body weight, and consequently die between 13-15 months of age (*23–25*). Mitochondrial dysfunction and inflammation are key features of aging (*26*), yet whether the innate immune system contributes to POLG-related disease phenotypes and premature aging of mutator mice is unknown.

Here, we report that mutator mice exhibit a hyper-inflammatory innate immune status that is driven by chronic engagement of the cGAS-STING-IFN-I axis. Persistent IFN-I signaling represses Nrf2 activity, which increases oxidative stress and aerobic glycolysis that potentiate inflammation and age-related pathology in these animals. Our findings indicate that IFN-I signaling is a key driver of innate immune rewiring and multi-organ pathology in mutator mice and provide a strong rationale for more broadly examining IFN-I dysregulation in mitochondrial diseases and aging.

## Results

### Innate immune hyper-responsiveness of POLG mutator mice is regulated by monocyte and neutrophil expansion

To begin to characterize innate immune alterations in the POLG mutator model of mtDNA disease, we employed a lipopolysaccharide (LPS)-induced endotoxemia model and monitored circulating cytokines and survival post challenge. Interestingly, both 6- and 12-month old mutator cohorts succumbed faster to intraperitoneal (i.p.) LPS challenge (Figure 1A and S1C). In-line with a more rapid mortality rate, we detected elevated levels of pro-inflammatory cytokines, chemokines, and type I interferons in the plasma of LPS-challenged mutator mice at all ages compared to WT littermates (Figure 1B, S1A and S1B), consistent with a prior report (*27*).

**Figure 1.**
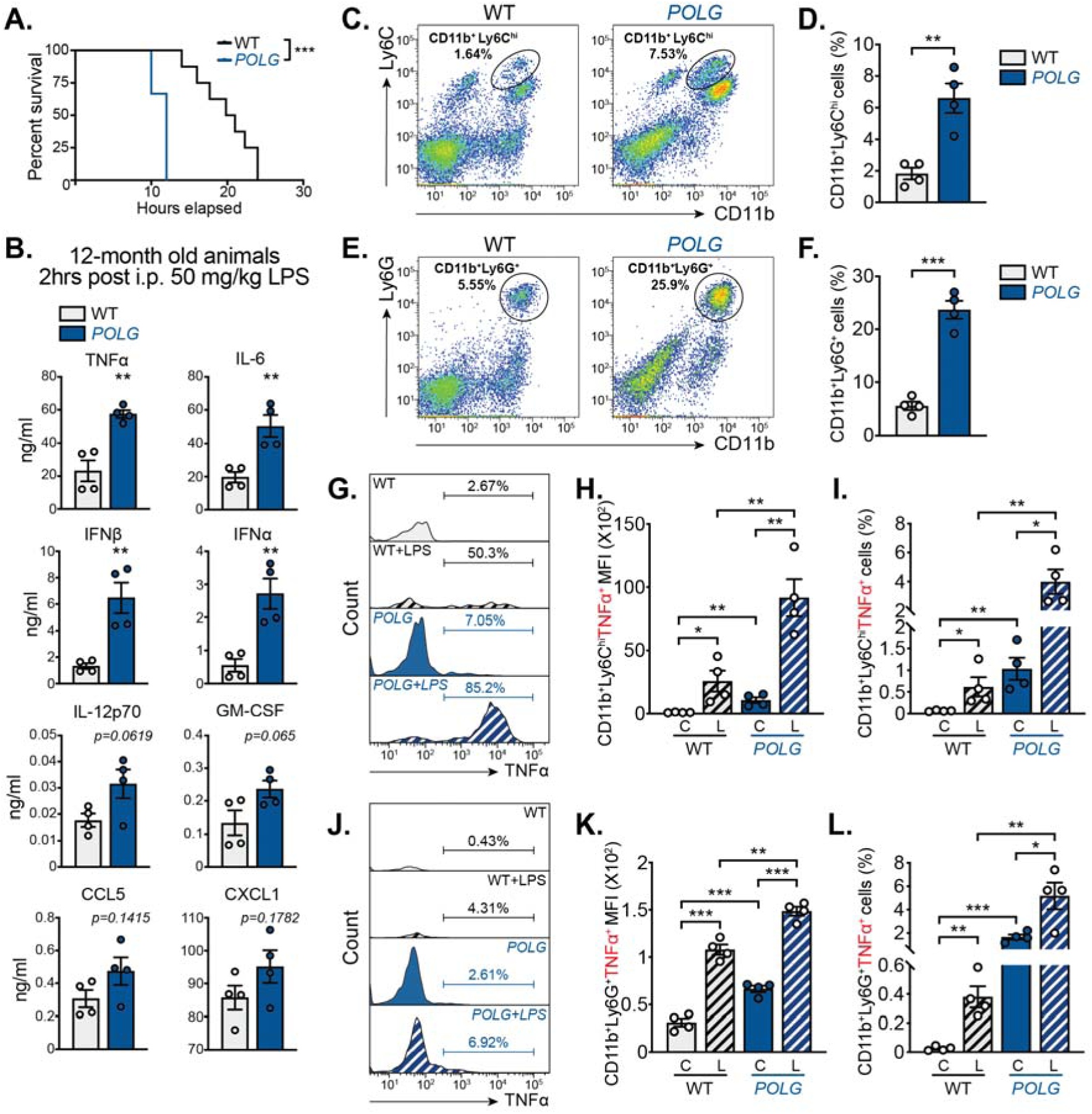
**POLG mutator mice exhibit a hyper-inflammatory phenotype to LPS challenge due to increased CD11b^+^ myeloid cells in the blood.** (A and B) 12-month old WT (n=8) and *POLG* (n=7) mice were challenged with LPS (50 mg/kg by i.p. injection). Kaplan-Meyer survival analysis was performed (A). Plasma cytokine profiles were determined by multi-analyte bead-based immunoassay on two biological duplicates and two technical duplicates per group (B). Log-rank (Mantel-Cox) test was used to compare percent survival between different groups. (C and D) CD11b^+^Ly6C^hi^ inflammatory monocyte population in whole blood from 12-month old WT and *POLG* mice was evaluated by flow cytometry. Pseudocolor plots are representative of 4 independent experiments (C) and quantification of the percentage of CD11b^+^Ly6C^hi^ cells is shown in (D). (E and F) CD11b^+^Ly6G^+^ blood neutrophil population in 12-month old WT and *POLG* mice was determined by flow cytometry. Pseudocolor plots are representative of 4 independent experiments (E) and quantification of the percentage of CD11b^+^Ly6G^+^ cells is shown in (F). (G-I) CD11b^+^Ly6C^hi^TNFα^+^ inflammatory monocyte population in unstimulated [C] or LPS challenged [L] whole blood from 12-month old WT and *POLG* mice was evaluated by flow cytometry. Histograms are representative of 4 independent experiments (G). Quantification of CD11b^+^Ly6C^hi^TNFα^+^ mean fluorescent intensity (MFI) is shown in (H), and the percentage of CD11b^+^Ly6C^hi^TNFα^+^ cells is shown in (I). (J-L) CD11b^+^Ly6G^+^TNFα^+^ neutrophil population in unstimulated [C] or LPS challenged [L] whole blood from 12-month old WT and *POLG* mice was evaluated by flow cytometry. Histograms are representative of 4 independent experiments (J). Quantification of CD11b^+^Ly6G^+^TNFα^+^ MFI is shown in (K), and the percentage of CD11b^+^Ly6G^+^TNFα^+^ neutrophils is shown in (L) . Unless stated, statistical significance was determined using unpaired Student’s t-tests. \**P* < 0.05; \*\**P* < 0.01; \*\*\**P* < 0.001. Error bars represent S.E.M.

As circulating leukocyte populations are key mediators of the cytokine storm that contributes to endotoxin shock, we next examined the blood cell composition of 12-month old cohorts. Although aged mutator mice exhibited B and T cell lymphopenia (Figure S1D and S1E) (*28, 29*), we noted a significant expansion of innate immune cell types, namely monocyte (CD11b^+^, Ly6C^hi^) and neutrophil (CD11b^+^, Ly6G^+^) populations, by flow cytometry (Figure 1C-1F). We also observed significant increases in steady-state tumor necrosis factor alpha (TNFα) and monocyte chemoattractant protein 1 (MCP-1/CCL2) in the plasma of aged mutators (Figure S1F). To next identify the sources of TNFα at baseline and after challenge, we employed intracellular staining and multiparameter flow cytometry. B and T cell populations in mutator mice did not express higher levels of intracellular TNFα before or after ex vivo LPS stimulation (Figure S1G). However, CD11b^+^ cells expressed significantly more TNFα and IL-6 after challenge (Figure 1G, 1J and S1H). Quantitation across multiple experiments revealed that CD11b^+^Ly6C^hi^ inflammatory monocytes in mutator blood possessed the greatest intracellular TNFα intensity, both at rest and after LPS stimulation (Figure 1G-1I). Although CD11b^+^Ly6G^+^ neutrophils were also more numerous in the mutator blood, their TNFα intensity was roughly 100-fold lower (Figure 1J-1L), indicating that inflammatory monocyte expansion is most likely responsible for the elevated levels plasma cytokines at rest and after Toll-like receptor 4 (TLR4) engagement. Similar expansion and elevated TNFα positivity was observed in monocyte and neutrophil lineage cells in the bone marrow of mutator mice (Figure S1I and S1J). Finally, oral infection of 9-10-month old cohorts with *Listeria monocytogenes* resulted in notably higher pro-inflammatory cytokine levels in mutator plasma at day 3 (Figure S1K), but less bacterial dissemination to the spleen and liver at day 5 post infection (Figure S1L). These results are consistent with prior findings that Ly6C^hi^ inflammatory monocytes predominantly control oral *Listeria* infection (*30*), and taken together, suggest that CD11b^+^ myeloid cell expansion and innate immune reprogramming drive systemic hyper-inflammatory responses in mutator mice.

### STING regulates enhanced IFN-I and pro-inflammatory responses in POLG mutator macrophages

To define the underlying signaling pathways that shape innate immune hyper-responsiveness in mutator mice, we performed RNA sequencing analysis of primary bone marrow-derived macrophages (BMDMs) and peritoneal macrophages (PerMacs) at rest and after LPS challenge. Pathway analysis of RNAseq datasets revealed significant elevations in interferon/IRF, JAK/STAT, and NF-κB signaling, as well as increased glycolytic metabolism and reactive oxygen and nitrogen species production in mutator macrophages (Figure S2A). Gene expression profiling of mutator macrophages revealed an enrichment of interferon-stimulated genes (ISG) (Figure 2A and 2B), which was confirmed by qRT-PCR analyses (Figure 2C, 2D, S2B and S2C). Mutator macrophages also displayed augmented pro-inflammatory cytokine RNA (Figure 2E and 2F) and protein levels (Figure 2G and 2H), agreeing with the intracellular cytometry data of Figure 1. Moreover, mutator macrophages produced higher levels of nitrite after LPS and IFNγ co-stimulation, confirming pathway analysis of RNAseq data (Figure 2I). Finally, interferon stimulatory DNA (ISD) transfection to directly engage cGAS-STING revealed elevated ISG and pro-inflammatory cytokine expression in mutator BMDMs (Figure 2J), demonstrating that POLG mutator macrophages are broadly hyper-responsive to multiple innate immune stimuli.

**Figure 2.**
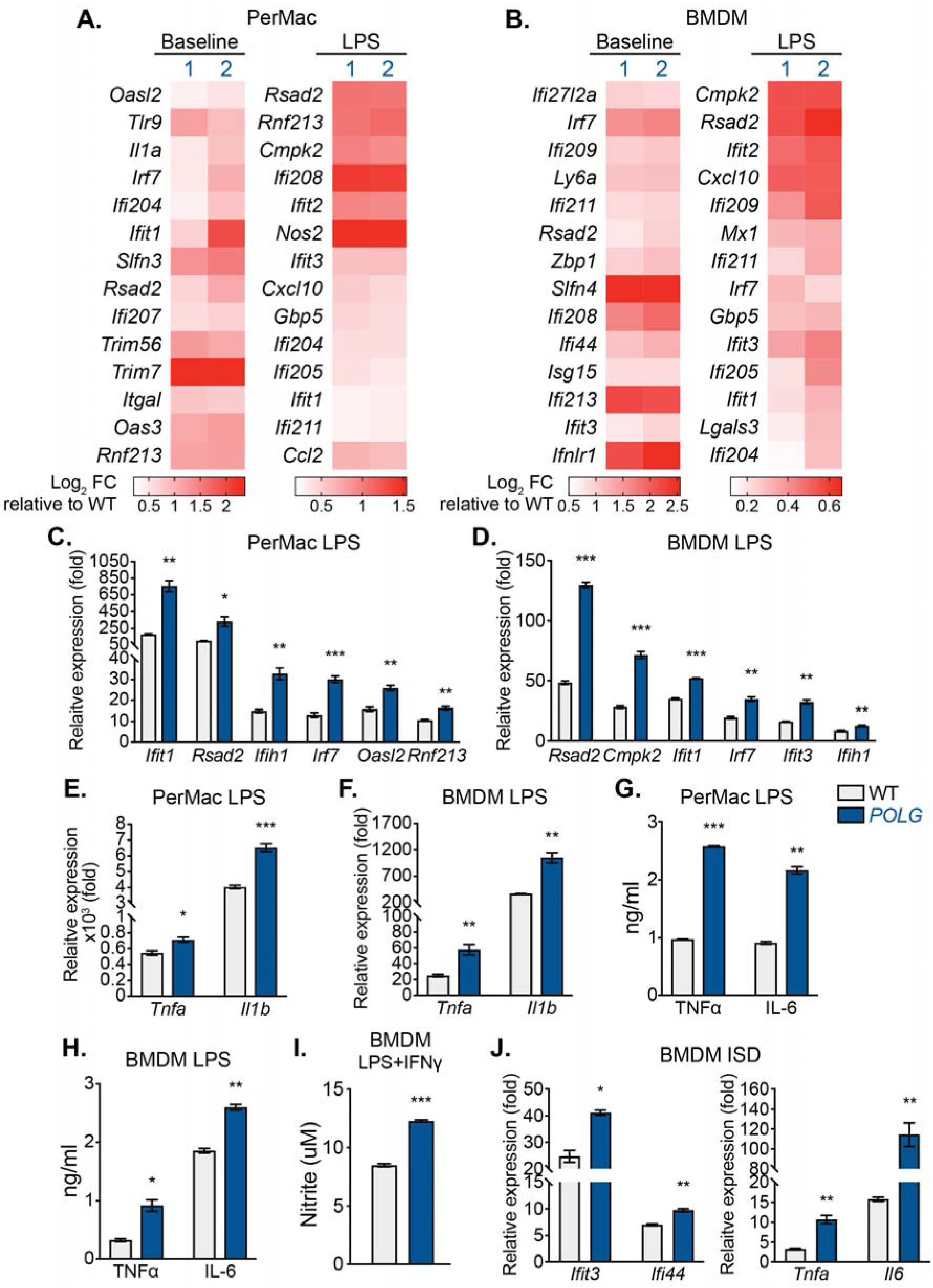
**POLG mutator macrophages exhibit enhanced IFN-I and pro-inflammatory responses after innate immune stimulation.** (A and B) Heatmap of RNAseq data displaying statistically significant (*P* < 0.05 in two biological replicates) differences in ISG expression between WT and *POLG* mutator peritoneal macrophages (PerMacs) (A) and bone marrow-derived macrophages (BMDMs) (B) post LPS challenge (200ng/ml for 6h). (C-F) qRT-PCR analysis of ISG and pro-inflammatory cytokine expression in WT and *POLG* PerMacs (C and E) and BMDM (D and F) after 4h or 6h of LPS stimulation. (G and H) Pro-inflammatory cytokine secretion in WT and *POLG* PerMacs (G) or BMDMs (H) after 4h or 6h of LPS stimulation. (I) Nitrite levels in WT and *POLG* BMDM after 17h of LPS (20ng/ml) +IFNγ (50ng/ml) treatment. (J) qRT-PCR analysis of ISGs and cytokines expression in WT and *POLG* BMDMs after 4h of 2ug/ml ISD transfection. Statistical significance was determined using unpaired Student’s t-tests. \**P* < 0.05; \*\**P* < 0.01; \*\*\**P* < 0.001. Error bars represent S.E.M.

STING is a key mediator of IFN-I and pro-inflammatory responses induced by cytosolic and extracellular mtDNA (*6, 7*), and ablation of STING can limit mtDNA-driven inflammation in mouse models of acute kidney injury and Parkinson’s disease (*31–33*). We observed that aged POLG mutator mice have significantly more circulating, cell-free mtDNA in the plasma, and we also noted that LPS-challenged mutator BMDMs liberate more mtDNA into the culture media (Figure S2D and S2E). We therefore reasoned that mtDNA instability and release in POLG mutator macrophages might drive constitutive IFN-I and elevated pro-inflammatory responses after LPS via the cGAS-STING pathway. Notably, we observed that macrophages from POLG mutators crossed onto a STING-deficient background had lower ISG expression (Figure S2F and S2G), as well as lower TNFα secretion (Figure S2H), when compared to STING-sufficient mutators. Taken together, our data suggest that mtDNA instability and release in POLG mutators engages STING, which potentiates macrophage activation to subsequent innate immune challenge.

### The cGAS-STING-IFN-I signaling axis regulates monocyte and neutrophil expansion and hyper-inflammatory responses in mutator mice

The early, inflammatory phase of septic shock is characterized by leukocyte activation and secretion of pro-inflammatory cytokines such as TNFα and IL-1β (*34*). IFN-I can exacerbate inflammation in response to TNFα (*35*) and contributes to mortality in LPS-induced sepsis (*36*). Because we noted increased IFN-I secretion after LPS challenge (Figure 1B) and more mtDNA was present in the plasma of aged POLG mutators (Figure S2D), we explored whether cGAS-STING signaling contributes to increased mortality after i.p. LPS challenge. Interestingly, we observed delayed LPS-induced mortality (Figure 3A) and markedly lower plasma cytokine levels in mutator mice lacking cGAS or STING (Figure 3B). In line with a role for IFN-I signaling in driving hyper-inflammatory responses to LPS, aged mutator mice lacking type I interferon receptor subunit 1 (*Ifnar^-/-^*) also had lower circulating levels of many cytokines compared to mutator mice alone (Figure 3B, rightmost columns).

**Figure 3.**
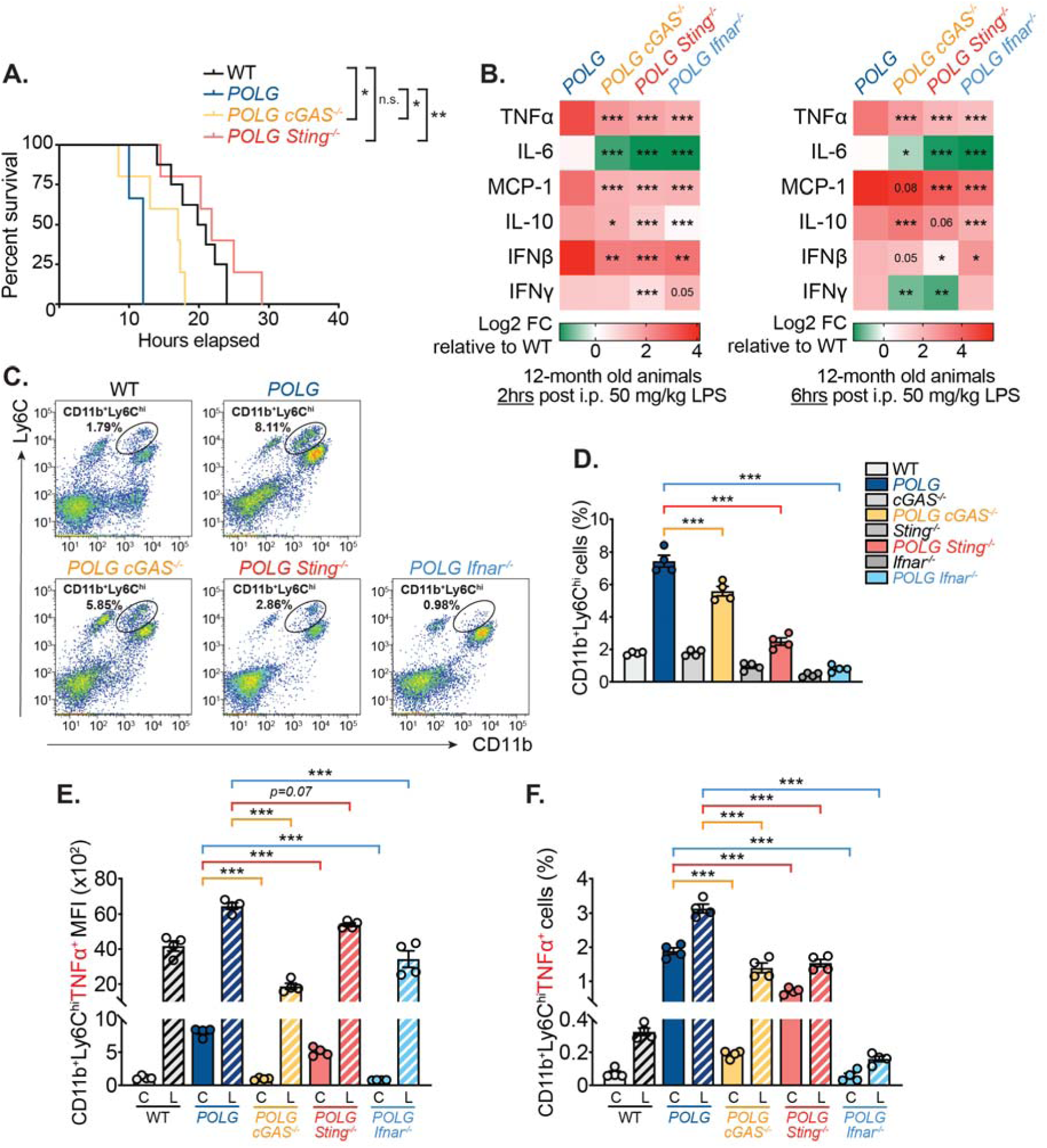
**The cGAS-STING-IFN-I signaling axis regulates inflammatory monocyte expansion and elevated cytokine secretion in POLG mutator mice.** (A and B) 12-month old WT, *POLG*, *POLG cGAS^-/-^*, *POLG Sting^-/-^* and *POLG Ifnar^-/-^* (n=5-8 per group) mice were i.p. injected with 50 mg/kg LPS. Kaplan-Meyer survival analysis was performed (A). Plasma was collected at indicated time points (n=6 at 2h and n=4 at 6h) and subjected to multi-analyte cytokine analysis (B). Statistical comparisons (B) were made against LPS injected *POLG* mice. Log-rank (Mantel-Cox) test was used to compare percent survival between different groups. (C and D) CD11b^+^Ly6C^hi^ inflammatory monocyte population in whole blood from 12-month old mice was evaluated by flow cytometry. Pseudocolor plots are representative of 4 independent experiments (C) and quantification of the percentage of CD11b^+^Ly6C^hi^ cells is shown in (D). (E and F) CD11b^+^Ly6C^hi^TNFα^+^ inflammatory monocyte population in unstimulated [C] or LPS challenged [L] whole blood from 12-month old cohorts was evaluated by flow cytometry. Quantification of CD11b^+^Ly6C^hi^TNFα^+^ mean fluorescent intensity (MFI) is shown in (E), and the percentage of CD11b^+^Ly6C^hi^TNFα^+^ cells is shown in (F). Unless stated, statistical significance was determined using unpaired Student’s t-tests or ANOVA. \**P* < 0.05; \*\**P* < 0.01; \*\*\**P* < 0.001. Error bars represent S.E.M.

IFN-I signaling is known to drive peripheral myeloid expansion in murine models of lupus and can modulate Ly6C^hi^ inflammatory monocyte recruitment in both infectious and sterile diseases (*37–41*). To next assess whether cGAS-STING-IFN-I activation governs increased peripheral myeloid expansion in mutator mice, we examined CD11b^+^Ly6C^hi^ monocyte and CD11b^+^Ly6G^+^ neutrophil populations in double mutant cohorts. Notably, CD11b^+^Ly6C^hi^ inflammatory monocyte numbers were progressively lower in cGAS-, STING-, and IFNAR-deficient mutators (Figure 3C and 3D), while blood neutrophil abundance was also decreased (Figure S3A and S3B). Moreover, both CD11b^+^Ly6C^hi^ monocytes (Figure 3E and 3F) and CD11b^+^Ly6G^+^ neutrophils (Figure S3C and S3D) from IFNAR-deficient mutators produced less TNFα after ex vivo LPS challenge. Additionally, TNFα^+^IL-1β^+^ double positive leukocytes were decreased in IFNAR-deficient mutator blood after LPS challenge (Figure S3E), and the elevated percentage of CD11b^+^Ly6C^hi^TNFα^+^ cells in LPS-stimulated mutator bone marrow was ablated by IFNAR knockout (Figure S3F). Collectively, these results suggest that sustained cGAS-STING-IFN-I signaling in POLG mutator mice promotes peripheral myeloid expansion and innate immune rewiring that potentiates systemic inflammatory responses and increases mortality to LPS challenge.

### Antioxidant and anti-inflammatory Nrf2 signaling is suppressed in POLG mutator macrophages

To next define the mechanisms underlying elevated ISG and pro-inflammatory cytokine expression in mutator monocytes and macrophages, we assayed key steps in NF-κB, IRF, and STAT1 activation during a 24-hour LPS time course. Although pathway analysis suggested potentiated signaling in mutator macrophages (Figure S2A), the kinetics of NF-κB, IRF, and STAT1 activation were nearly identical between WT and mutator BMDMs following LPS treatment (Figure S4A). However, in agreement with RNAseq data implicating elevated glycolytic metabolism, we observed a marked increase in the extracellular acidification rate (ECAR) and decreased basal and maximal oxygen consumption rates (OCR) in mutator BMDMs (Figure S4B and S4C). This suggested metabolic reprogramming away from mitochondrial respiration to glycolysis for enhanced inflammatory M1 macrophage activation. A recent study reported that the nuclear factor erythroid 2-related factor 2 (Nrf2) transcription factor interferes with LPS-induced cytokine gene expression and is therefore a major transcriptional repressor of M1 macrophage polarization and inflammation (*42*). Our RNA profiling indicated that a significant number of Nrf2-target genes were transcriptionally repressed in mutator macrophages (Figure S2A and 4A). In agreement, analysis of endogenous Nrf2 by quantitative fluorescent microscopy revealed markedly lower nuclear translocation in LPS-stimulated POLG mutator macrophages (Figure 4B-4D).

**Figure 4.**
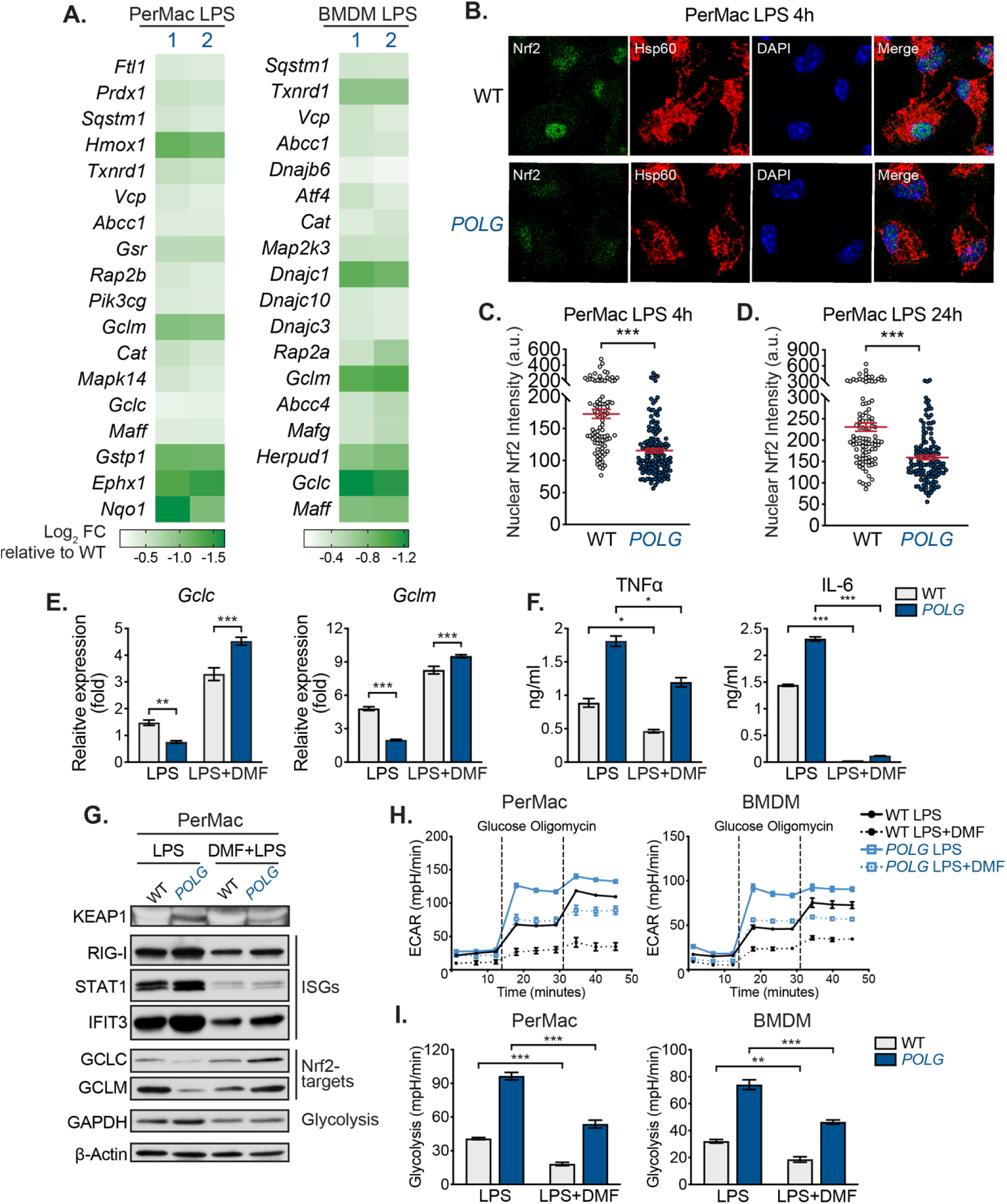
**Nrf2 suppression contributes to the hyper-inflammatory phenotype of POLG mutator macrophages.** (A) Heatmap of RNAseq data displaying statistically significant (*P* < 0.05 in two biological replicates) differences in Nrf2 target gene expression between WT and *POLG* mutator PerMacs BMDMs post LPS challenge (200ng/ml for 6h). (B) Representative confocal microscopy images of LPS treated PerMacs stained with anti-Nrf2 and -Hsp60 antibodies and DAPI. (C and D) Quantification of nuclear Nrf2 staining intensity in WT and *POLG* PerMacs 4h (C) or 24h (D) post LPS stimulation. a.u., arbitrary unit. (E) qRT-PCR analysis of Nrf2 target gene expression in WT and *POLG* PerMacs post LPS or LPS+DMF treatment. (F) Pro-inflammatory cytokine secretion by WT and *POLG* PerMacs after LPS or LPS+DMF treatment. (G) Protein expression in WT and *POLG* PerMacs post LPS or LPS+DMF treatment. (H and I) Seahorse ECAR analysis of WT and *POLG* PerMacs and BMDMs post LPS or LPS+DMF exposure (H). Glycolysis rate was calculated and plotted in (I). Statistical significance was determined using unpaired Student’s t-tests. \**P* < 0.05; \*\**P* < 0.01; \*\*\**P* < 0.001. Error bars represent S.E.M.

To next explore whether Nrf2 suppression contributes to hyper-inflammatory phenotypes in mutator macrophages, we employed two pharmacological approaches to boost Nrf2 activity. Itaconate functions as an anti-inflammatory metabolite by alkylating Kelch-like ECH-associated protein 1 (KEAP1) (*43*), which normally associates with and promotes the degradation of Nrf2. Itaconate-mediated alkylation of key KEAP1 cysteine residues allows newly synthesized Nrf2 to accumulate, translocate to the nucleus, and activate a transcriptional antioxidant and anti-inflammatory program (*43, 44*). Supplementing POLG mutator PerMacs with 4-octyl-itaconate (4OI) (a cell-permeable itaconate derivative) restored the expression of Nrf2 target genes (Figure S4D). In addition, 4OI markedly reduced the elevated secretion of pro-inflammatory cytokines in LPS-challenged mutator macrophages (Figure S4E). Treatment of macrophages with a second Nrf2 stabilizer, dimethyl fumarate (DMF), also restored Nrf2 target gene expression (Figure 4E and 4G), suppressed inflammatory cytokine production (Figure 4F), and reduced ISGs in mutator macrophages (Figure 4G). Interestingly, we noted that KEAP1 protein was markedly elevated in mutator PerMacs after LPS, while being nearly undetectable in WT cells (Figure 4G). DMF also limits aerobic glycolysis and lowers cytokine secretion from activated macrophages by inactivating GAPDH (*45*). Accordingly, DMF reduced the elevated GAPDH protein expression in PerMacs from mutator mice (Figure 4G), while also reducing aerobic glycolysis in both WT and mutator macrophages (Figure 4H and 4I). Together, these data indicate that Nrf2 suppression and augmented aerobic glycolysis shift POLG mutator macrophages toward a more pro-inflammatory state.

### IFN-I signaling represses Nrf2 activity and drives pro-inflammatory metabolic phenotypes in POLG mutator macrophages

Nrf2 has been recently reported to inhibit STING-dependent IFN-I responses (*46–48*), and IFN-I signaling can limit microbial clearance by dysregulating Nrf2 target gene expression (*49, 50*). We therefore utilized IFNAR-deficient cohorts to examine whether hyperactive IFN-I signaling contributes to Nrf2 suppression in mutator macrophages. We first confirmed that IFNAR depletion ablated elevations in LPS-induced ISG expression in mutator macrophages (Figure 5A). Next, we utilized quantitative fluorescent microscopy and observed that *Ifnar^-/-^* mutator macrophages displayed marked restoration of Nrf2 nuclear translocation after LPS challenge (Figure 5B-5D and S5A-5C). RNA and protein profiling of Nrf2 targets further confirmed that ablation of IFNAR was sufficient to restore, or even hyperactivate, Nrf2 signaling in *Ifnar^-/-^* mutator macrophages (Figure 5E, 5F and S5D). In addition, the secretion of TNFα from *Ifnar^-/-^* mutator PerMacs was lowered (Figure S5E), further confirming our intracellular cytokine staining results (Figure 3 and S3).

**Figure 5.**
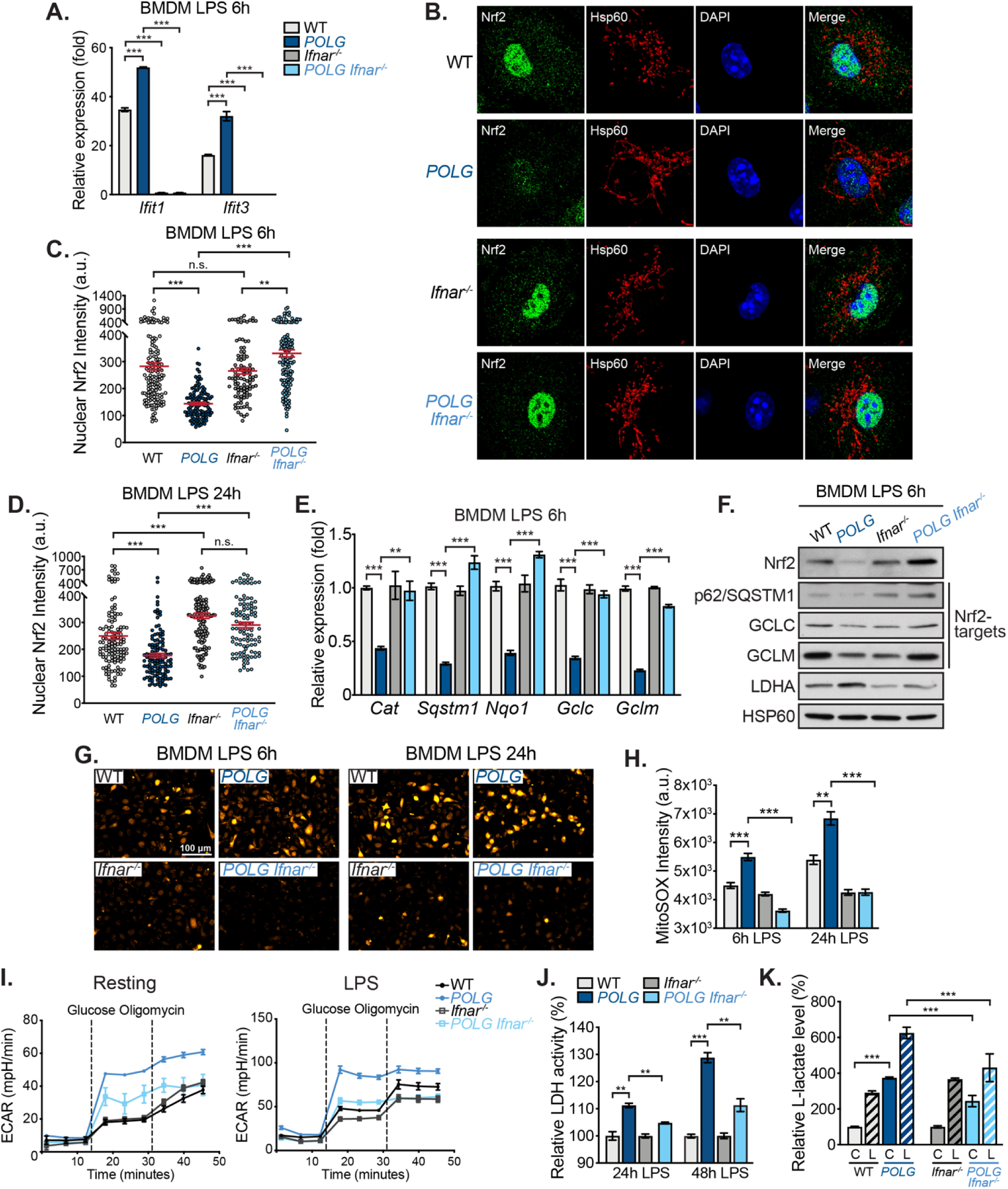
**Elevated IFN-I signaling represses Nrf2 activity and drives pro-inflammatory metabolic phenotypes in POLG mutator macrophages.** (A) qRT-PCR analysis of ISG expression in BMDMs after LPS challenge. (B) Representative confocal microscopy images of LPS treated BMDMs stained with anti-Nrf2 and -Hsp60 antibodies and DAPI. (C and D) Quantification of nuclear Nrf2 staining intensity in BMDMs 6h (C) or 24h (D) after LPS exposure. a.u., arbitrary unit. (E) qRT-PCR analysis of Nrf2 target gene expression in BMDMs post LPS exposure. *POLG* BMDMs were normalized to WT BMDMs, POLG *Ifnar^-/-^* BMDMs were normalized to *Ifnar^-/-^* BMDMs. (F) Protein expression of Nrf2 targets and LDHA in BMDMs after 6h LPS exposure. (G) Representative microscopy images of LPS treated BMDMs stained with MitoSOX. (H) Quantification of MFI of MitoSOX staining in LPS treated BMDMs. (I) Seahorse ECAR analysis of BMDMs with or without overnight LPS (10ng/ml) challenge. (J) Relative LDH activity in BMDMs 24h and 48h post LPS exposure (normalized to WT). (K) Extracellular L-Lactate level in culture media of resting [C] or LPS exposed [L] BMDMs. WT and *POLG* were normalized to WT resting BMDMs, *Ifnar^-/-^* and POLG *Ifnar^-/-^* were normalized to *Ifnar^-/-^* resting BMDMs. Statistical significance was determined using unpaired Student’s t-tests or ANOVA. \**P* < 0.05; \*\**P* < 0.01; \*\*\**P* < 0.001. Error bars represent S.E.M.

In line with a role for Nrf2 antioxidant programs in regulating both mitochondrial and cytosolic ROS (*51*), we detected significantly higher levels of mitochondrial superoxide and total cellular ROS in mutator BMDMs after LPS induction (Figure 5G, 5H, S5F and S5G). IFNAR depletion lowered ROS in POLG WT macrophages and dramatically reduced ROS in POLG mutator BMDMs. As we previously observed that POLG mutator macrophages exhibit elevated KEAP1 protein after LPS stimulation (Figure 4G), we next explored whether IFN-I directly affects KEAP1 abundance. Interestingly, we observed that treatment of BMDMs with recombinant mIFNβ augmented KEAP1 levels, leading to a concomitant reduction in Nrf2 abundance (Figure S5H). Overall, these results highlight that IFN-I-mediated repression of Nrf2, via increased KEAP1 expression, lowers antioxidant capacity and elevates pro-inflammatory ROS in mutator macrophages.

IFN-I signaling can shift innate immune cell metabolism from OXPHOS to aerobic glycolysis due to induced breaks in the TCA cycle (*52, 53*). Accordingly, we found that both acute and chronic IFNβ treatment slightly elevated ECAR in WT BMDMs, but dramatically increased ECAR in mutator macrophages (Figure S5I). In contrast, knockout of IFNAR markedly reduced ECAR in resting and LPS-stimulated in mutator macrophages, while having little to no effect on WT macrophages (Figure 5I). Consistent with the changes in ECAR, mutator macrophages had higher LDH expression level and activity, and thus produced more L-lactate than WT cells. Ablation of IFN-I signaling in POLG BMDMs largely reversed elevated LDH expression (Figure 5F) and activity (Figure 5J) after LPS challenge, while also blunting L-lactate levels (Figure 5K). Taken together, our results suggest that blocking IFN-I signaling restores Nrf2, lowers oxidative stress and aerobic glycolysis, and blunts the hyper-inflammatory profile in POLG mutator macrophages.

### Blocking IFN-I signaling restores Nrf2 activity, limits oxidative stress, and lowers aerobic glycolysis in aged POLG mutator mice

Several studies have linked oxidative stress to tissue dysfunction and premature aging phenotypes in POLG mutator mice (*27, 54–57*). Therefore, we next explored dysregulation in IFN-I-Nrf2 crosstalk in the heart, liver, and kidney of 12-month old animals. Transcript and protein profiling revealed a significant increase in ISGs, similar to that observed in POLG macrophages, which was dependent on intact IFNAR signaling (Figure S6A and S6B). In addition, the expression of Nrf2 and Nrf2-regulated antioxidant genes (*Nqo1, Gclc, Gclm, Sqstm1/p62*) was markedly lower in mutator tissues compared to age-matched WT littermates, but was rescued in *Ifnar^-/-^* mutator tissues (Figure S6C and S6D). Aconitase is widely recognized as a sensitive and specific target of ROS, and aconitase inactivation is a surrogate marker of oxidative stress. Therefore, to determine whether IFN-I-mediated Nrf2 suppression increases oxidative stress in mutator mice, we analyzed the enzymatic activity of aconitase in various tissues of WT, mutator, *Ifnar^-/-^*, and *Ifnar^-/-^* mutator littermate cohorts. Heart, liver, and kidney extracts from mutator mice showed significantly lower aconitase activity, indicative of oxidative stress; however, IFNAR deficiency largely restored aconitase activity in mutator tissues, likely via boosting Nrf2-regulated antioxidant activity (Figure S6E).

Our prior observations in macrophages and other reports indicated imbalances in glycolytic metabolism and oxidative phosphorylation in mutator tissues (*58*), thus we next examined expression of enzymes in glycolysis, the TCA cycle, and OXPHOS (Figure S6F). GAPDH and LDHA were significantly elevated in aged POLG mutator tissues (Figure S6F, top panel); however, *Ifnar^-/-^* mutator cohorts exhibited decreased glycolytic enzyme expression and decreased L-lactate accumulation in the plasma (Figure S6F and S6G). In addition, several TCA cycle enzymes were upregulated in mutator heart and liver homogenates, which may reflect a compensatory response to elevated aerobic glycolysis and OXPHOS deficiency. Consistent with a role for IFN-I-mediated metabolic rewiring, TCA enzymes were largely restored to WT levels in *Ifnar^-/-^* mutator tissues (Figure S6F, middle panel). Finally, *Ifnar^-/-^* mutator cohorts exhibited modestly higher levels of some OXPHOS subunits, namely mt-CO1 and NDUFB8 in the liver, compared to IFNAR-sufficient mutators (Figure S6F, bottom panel). Collectively, our tissue analyses mirror the key findings from macrophage studies, and thus define chronic IFN-I responses as critical regulators of Nrf2 suppression and metabolic rewiring in POLG mutator mice.

### Ablation of IFN-I signaling improves healthspan and extends lifespan of POLG mutator mice

Age-related multi-tissue pathology, including alopecia, kyphosis, anemia, and dilated cardiomyopathy, has been noted in numerous reports on POLG mutator mice (*23, 24, 28, 55, 59*). As both elevated IFN-I signaling and Nrf2 inhibition are implicated in cardiomyopathy and anemia in animal models and human patients (*60–63*), we reasoned that sustained imbalances in IFN-I and Nrf2 signaling may contribute to the aging-related phenotypes of mutator mice. Using transthoracic echocardiography, we confirmed that 9-10-month old mutator mice exhibited dilated cardiomyopathy by demonstrating left ventricle dilation (wider left ventricular inner diameter at diastole LVID;d) and markedly reduced systolic function (decreased left ventricular ejection fraction, LVEF) (Figure 6A and 6B). Other dilated cardiomyopathy phenotypes included decreased interventricular septum thickness at end-systole (IVS;s), increased left ventricular internal dimension at end-systole (LVID;s) and elevated left ventricular volume at end-systole (Vol;s) (Figure S7A and S7B). Remarkably, cardiac dilation and LVEF were improved in IFNAR-deficient mutator cohorts (Figure 6A, 6B, S7A and S7B). In addition, histological analyses revealed increased mean myofiber width and more infiltrating immune cells in mutator hearts, which were both significantly reduced in IFNAR-deficient mutator sections (Figure 6C and 6D). Finally, we noted a shift in the cardiac myosin heavy chain isoform from *Myh6* toward *Myh7* in mutator heart homogenates, which is a well-appreciated marker of cardiac hypertrophy, injury, and stress. Consistent with our echocardiographic and histological measurements, IFNAR-deficient mutator hearts exhibited increased *Myh6* and reduced *Myh7* expression, indicative of lower cardiomyopathy (Figure S7C).

**Figure 6.**
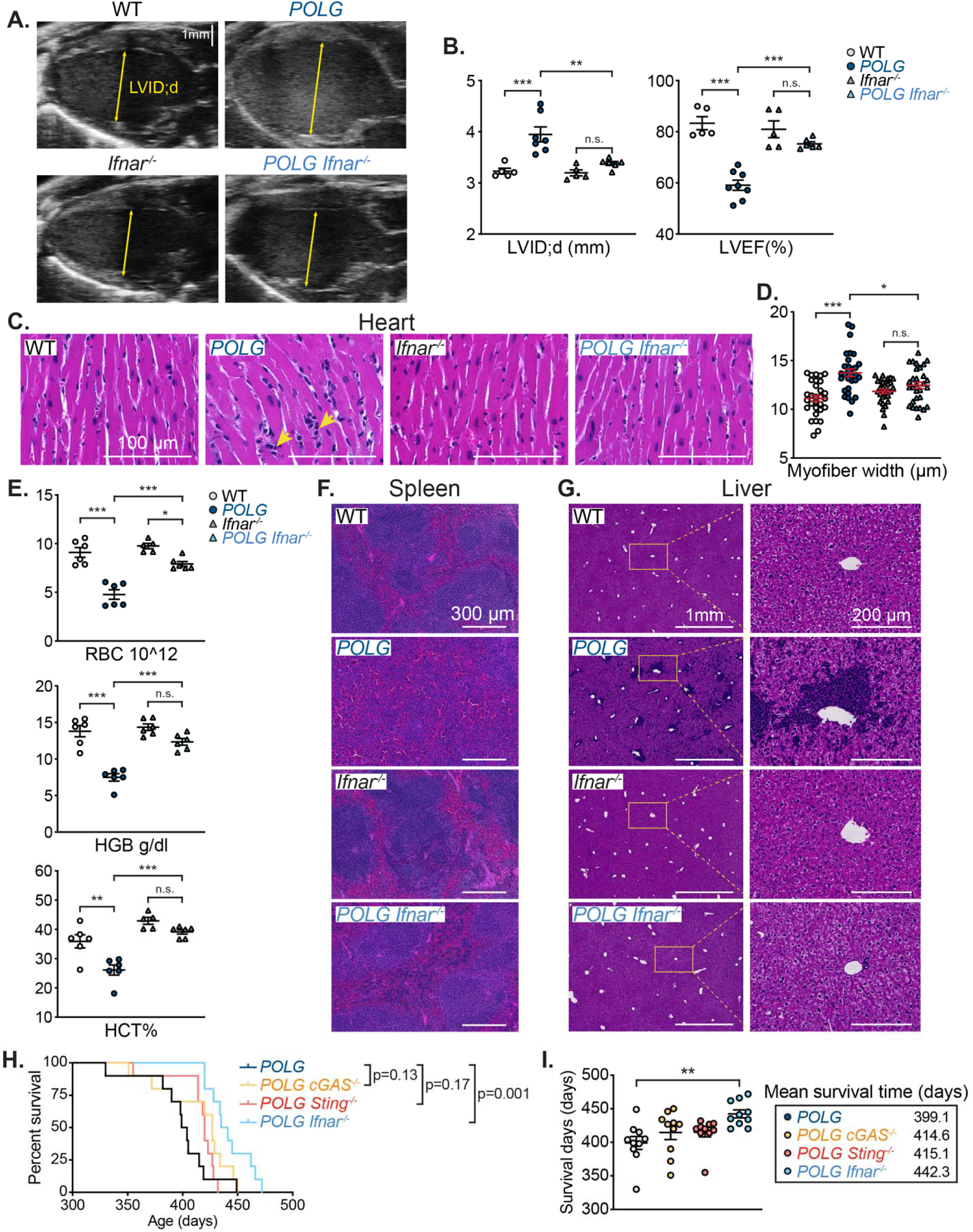
**Ablation of IFN-I signaling lessens multi-organ pathology and extends lifespan in POLG mutator mice.** (A) Representative B-mode echocardiogram images of 9-10-month old WT, *POLG*, *Ifnar^-/-^* and *POLG Ifnar^-/-^* mouse hearts. (B) Left ventricular internal dimension at end-diastole (LVID;d) and left ventricular ejection fraction (LVEF) calculated from M-mode or B-mode images using Vevo Lab software. n=5-8 animals per genotype. (C and D) Representative H&E staining of heart sections from WT, *POLG*, *Ifnar^-/-^* and *POLG Ifnar^-/-^* mice (C) and quantification of cardiomyocyte width (D). Yellow arrows indicate infiltrating immune cells. 5 myocytes per section and 6 animals per genotype were quantified in a blinded fashion (D). (E) Red blood cell counts (RBC), hemoglobin concentration (HGB) and hematocrit (HCT) were measured in WT, *POLG*, *Ifnar^-/-^* and *POLG Ifnar^-/-^* mouse whole blood using the HM5 Hematology Analyzer. n=6 animals per genotype. (F) Representative H&E staining showing white pulp and red pulp organization in WT, *POLG*, *Ifnar^-/-^* and *POLG Ifnar^-/-^* mouse spleens. (G) Representative H&E stained liver sections from WT, *POLG*, *Ifnar^-/-^* and *POLG Ifnar^-/-^* cohorts. (H and I) Percent survival (H) and survival time (I) of *POLG*, *POLG cGAS^-/-^*, *POLG Sting^-/-^* and *POLG Ifnar^-/-^* mice. Log-rank (Mantel-Cox) test was used to compare percent survival between different groups. n=10 animals per genotype. Unless stated, statistical significance was determined using ANOVA.\**P* < 0.05; \*\**P* < 0.01; \*\*\**P* < 0.001. Error bars represent S.E.M.

POLG mice develop progressive, ultimately fatal megaloblastic anemia due to hematopoietic stem cell deficits, impaired erythrocyte maturation, and increased erythrocyte destruction by splenic macrophages (*28, 64, 65*). Remarkably, we found that *Ifnar^-/-^* mutator blood contains increased cellularity (Figure S7D), as well as higher red blood cell (RBC) counts, hemoglobin (HBG) concentration, and hematocrit (HCT) levels compared to IFNAR-sufficient mutators (Figure 6E). Moreover, ablation of IFNAR-signaling significantly restored splenic architecture in POLG mutator mice and increased white pulp to red pulp abundance (Figure 6F). We also observed widespread extramedullary hematopoiesis in the livers of mutator mice, which was largely ablated in age-matched *Ifnar^-/-^* mutator cohorts (Figure 6G), indicating that persistent IFNAR signaling in POLG proofreading-deficient mice contributes to the progressive and lethal anemia observed. Finally, we noted that loss of cGAS-STING-IFN-I signaling generally improved body condition in 12-month old mutator mice, as indicated by less kyphosis of the spine and reduced hair loss (Figure S7E). Remarkably, *Ifnar^-/-^* mutator cohorts exhibited a 10% extension in mean lifespan over IFNAR-sufficient mutator mice (Figure 6H and 6I). Altogether, our data suggest that innate immune reprogramming and chronic IFN-I signaling contribute to age-related anemia and cardiomyopathy, negatively impacting both healthspan and lifespan of POLG mutator mice (Figure S8).

## Discussion

Recent clinical case reports suggest that patients with MDs experience recurrent infections and develop sepsis and/or SIRS at significantly elevated rates relative to the general population (*14, 15, 66*). In fact, one study reported that sepsis is the most frequent cause of early death in pediatric MD patients (*13*). The innate immune system is the first line of defense against invading microbes and is an important driver of hyper-inflammatory responses in sepsis and SIRS. However, there is a paucity of information about innate immune system function and/or cellular composition in patients with MDs or murine models. Our work begins to address this critical knowledge gap in a mouse model of POLG-related MD. We have uncovered that POLG mutator mice are highly susceptible to lethal endotoxin shock owing to a dramatic increase in circulating, myeloid-derived innate immune cell populations and elevated levels of pro-inflammatory cytokines and IFN-I in the plasma after i.p. LPS challenge. Our data suggest that CD11b^+^ myeloid cell expansion, especially the CD11b^+^Ly6C^hi^ inflammatory monocyte population, is a key driver of augmented cytokine secretion in mutator mice, as intracellular TNFα levels in CD11b^+^Ly6C^hi^ mutator monocytes are elevated 4-fold compared to WTs. Although mutator mice do not directly model any particular human MD, CD11b^+^ myeloid cell expansion in the bone marrow and spleen has also been observed in the *Ndufs4^-/-^* model of Leigh Syndrome (*67, 68*). Therefore, future studies in other mouse models and patients are warranted to determine the degree to which monocyte and neutrophil expansion/polarization occurs in MDs. Monitoring circulating MCP-1/CCL2, which is basally elevated in mutator mice and is a predictive biomarker for sepsis, may also aid in identifying MD patients at risk for developing life-threatening systemic inflammation after infection (*69, 70*).

Mounting evidence suggests that the IFN-I-driven expansion of inflammatory monocytes is deleterious in autoimmune and inflammatory disorders such as lupus and pneumonia (*37, 38, 40, 71*). Elevated IFN-I signaling during chronic pattern recognition receptor (PRR) stimulation drives hematopoietic stem cell (HSC) skewing toward granulocyte/macrophage progenitors (GMP), leading to emergency myelopoiesis in the bone marrow and peripheral myeloid expansion (*37, 72, 73*). Myeloid lineage skewing in the bone marrow of mutator mice has been previously reported, although the mechanisms underlying this phenotype have remained unclear (*74*). Remarkably, the expansion of Ly6C^hi^ inflammatory monocytes and Ly6G^+^ neutrophils, systemic cytokine levels, and time to death after lethal LPS challenge is significantly reduced in mutator mice lacking cGAS, STING, or IFNAR. We therefore propose that mtDNA instability and release in POLG mutators triggers cGAS-STING-dependent IFN-I priming, which not only drives HSC skewing and peripheral myeloid expansion, but also potentiates responsiveness of circulating monocytes and neutrophils to innate immune stimuli. In addition to well-characterized mtDNA instability, a recent study documented dNTP pool imbalance and nuclear genome instability in mutator mice (*75*). Future work should therefore clarify the cellular sources of mtDNA and mechanisms of release, while also examining the contribution of nuclear DNA damage to elevated innate immune and IFN-I signaling in POLG mice.

We have found that mutator macrophages are broadly hyper-responsive to pattern recognition receptor agonists, although unexpectedly, the signaling kinetics of NF-κB, IRF, and STAT1 are nearly identical between WT and mutator BMDMs following LPS treatment. Instead, our data indicate that the hyper-inflammatory phenotype of mutator macrophages results from enhanced M1 polarization due to IFN-I dependent Nrf2 suppression and metabolic rewiring. Nrf2 can directly inhibit LPS-induced cytokine gene expression, and loss of Nrf2 activity is linked to mitochondrial dysfunction and increased glycolysis (*42, 76*). Compared to cells from age-matched mutator cohorts, IFNAR-deficient mutator macrophages display higher Nrf2 levels and target gene expression, while also generating less ROS and pro-inflammatory cytokines after LPS challenge. This dramatic restoration in Nrf2 activity in *Ifnar^-/-^* mutators is likely due to a lessening of IFN-I- and IL-10-mediated TCA cycle breaks (i.e. reduced aconitase and isocitrate dehydrogenase activity), which lower carbon flux toward itaconate (*53*). However, consistent with findings in fibroblasts and induced pluripotent stem cells (*56, 58*), POLG mutator BMDMs also exhibit elevated ECAR and decreased OCR at rest. Thus, the restoration of Nrf2 activity in *Ifnar^-/-^* mutator macrophages may also be due to reduced aerobic glycolysis, which directs pyruvate away from TCA metabolism toward lactate generation. Consistent with this notion, loss of IFN-I signaling markedly blunts ECAR and lactate levels in *Ifnar^-/-^* mutator macrophages, and also lowers plasma lactate concentrations in aged mice. In sum, our results support a model whereby mtDNA mutagenesis in POLG mutator macrophages potentiates IFN-I signaling, which enhances Keap1 to destabilize Nrf2, while also elevating LDHA levels and activity to potentiate aerobic glycolysis.

Our research has also uncovered that elevated IFN-I responses and reduced Nrf2 activity extend to multiple organs of aged mutator mice. Genetic ablation of IFNAR is sufficient to blunt potentiated ISG expression and markedly increase Nrf2 target gene abundance in the heart, liver, and kidney of aged cohorts. ROS and oxidative stress have been linked to organ dysfunction in aged POLG mice, with antioxidant therapy providing some benefits to overall healthspan (*27, 54, 57, 64*). We found that oxidative stress is significantly lower in *Ifnar^-/-^* mutator organs compared to IFNAR-sufficient cohorts, suggesting that IFN-I signaling triggered by mtDNA mutagenesis and instability inhibits Nrf2-mediated antioxidant responses in vivo. Interestingly, Nrf2-null mice display a spectrum of pathology that overlaps with POLG mutator mice, namely anemia, splenomegaly, cardiomyopathy, and increased susceptibility to lethal septic shock (*63, 77, 78*). Moreover, reduced Nrf2 activity is linked to numerous aging-related diseases (*79–81*), and Nrf2 repression was recently uncovered as a novel driver of oxidative stress and premature aging in Hutchinson-Gilford progeria syndrome (HGPS) (*82*). Elevated IFN-I responses have also been observed in HGPS and other progeroid syndromes, which share some overlapping phenotypes with mtDNA mutator mice (*83–85*). Strikingly, we found that *Ifnar^-/-^* mutator cohorts exhibit an extension in mean survival time of roughly forty days and an overall improvement in body condition compared to IFNAR-sufficient mutators, documenting that perturbed IFN-I-Nrf2 crosstalk is an unappreciated molecular mechanism contributing to premature aging in mutator mice.

Cardiac and hematologic analyses indicate that blockade of IFNAR signaling during aging yields striking improvements in cardiomyopathy and anemia-related phenotypes in mutator mice. Mitochondrial ROS contributes to cardiomyopathy in mutator mice (*55*), and our genetic and echocardiographic data suggest that shifting the balance from IFN-I toward Nrf2 activity lowers cardiac oxidative stress and improves cardiac function in aged mutator animals. However, additional IFN-I-dependent inflammatory and metabolic processes are likely dysregulated in aged mutator hearts. Future research on the mutator strains described here should yield new insight into roles for IFN-I dysregulation in both aging- and MD-related cardiomyopathy. Elevated IFN-I signaling has also been linked to anemia in aging and in various human diseases and animal models (*61, 73, 86*). A recent report characterized a unique subset of splenic hemophagocytes that differentiate from IFN-I expanded Ly6C^hi^ monocytes, which are responsible for anemia in a model of macrophage activation syndrome and lupus (*37, 73*). As we observed the IFN-I- dependent elevation of Ly6C^hi^ monocytes in mutator mice, it is likely that the restoration of peripheral erythrocyte numbers in *Ifnar^-/-^* mutator blood is results from less GMP skewing in the bone marrow and reduced destruction of erythrocytes by inflammatory hemophagocytes. Anemia is the most frequent hematological abnormality observed in patients with MD, and the presence of anemia negatively influences survival in patients with POLG-related disease (*87, 88*). Our work provides a strong rationale for translational research to explore whether IFN-I-Nrf2 signaling imbalances potentiate anemia in elderly/frail populations and patients with POLG-related MD.

In conclusion, we report that mtDNA mutator mice exhibit a hyper-inflammatory innate immune status that is driven by chronic IFN-I priming and Nrf2 repression. Our work constitutes the first detailed characterization of innate immune rewiring in a model of POLG-related disease and may provide a mechanistic framework for understanding why some MD patients are more prone to developing sepsis and SIRS following infection. Moreover, we have made the novel observation that innate immune dysregulation and IFN-I-mediated inflammaging contribute to several progeroid phenotypes in mutator mice. Therapeutic approaches aimed at rebalancing IFN-I-Nrf2 signaling may therefore represent a promising target for limiting runaway inflammation, combatting anemia, and improving overall healthspan in progeroid syndromes, mitochondrial diseases, and aging.

## Materials and Methods

### Mouse strains

C57BL/6J, *POLG^D257A/D257A^* mutator, *cGAS^-/-^*, *Sting^-/-^* (*Tmem173^gt^)* and *Ifnar1^-/-^* mice were purchased from The Jackson Laboratory and bred and maintained in multiple vivaria at Texas A&M University. *POLG^D257A/+^* breeder pairs used to generate *POLG^+/+^* and *POLG^D257A/D257A^* experimental mice (and *cGAS*-, *Sting*- and *Ifnar*-null intercrosses) were obtained from male *POLG^D257A/+^* to female C57BL/6J (or *cGAS*-, *Sting*- and *Ifnar*-null on a pure C57BL/6J background) crosses. All animal experiments were approved by the Institutional Animal Care and Use Committee (IACUC) at Texas A&M University.

### Antibodies and reagents

Anti-IFIT3 was a gift from G. Sen at Cleveland Clinic and anti-VIPERIN was a gift from P. Cresswell at Yale School of Medicine. Commercially obtained antibodies and reagents include: antibodies for immunoblotting: rabbit anti-TBK1 (3504), -p-TBK1 (5483), -STAT1 (9172), -p-STAT1 (7649), -RIG-I (4200), -p-IκBα (2859), -IκBξ (93726), -IRF1 (8478), -IRG1 (17805), - NRF2 (12721), -KEAP1 (8047) (Cell Signaling Technology); rabbit anti-LDHA (19987-1-AP), - p62/SQSTM1 (18420-1-AP), -IκBα (10268-1-AP), mouse anti-GAPDH (600004-1), -β-Actin (66009-1) (Proteintech); mouse anti-OXPHOS (ab110413), rabbit anti-GCLC (ab207777), -PDH (ab110333) (abcam); mouse anti-ACO2 (MA1-029), rabbit anti-GCLM (MA5-32783) (Invitrogen); goat anti-HSP60 (N-20) (Santa Cruz Biotechnology); rabbit-OGDH (HPA020347) (Sigma); antibodies and reagents used for flow cytometry included: PE/Cy7 anti-mouse TNF-α Antibody (506324), PerCP/Cy5.5 anti-mouse Ly-6C Antibody (128012), FITC anti-mouse Ly-6C Antibody (128006), PerCP/Cy5.5 anti-mouse IL-12/IL-23 p40 (505211) (BioLegend); Purified Anti-Mouse CD16 / CD32 (2.4G2) (70-0161), violetFluor™ 450 Anti-Mouse Ly-6G (1A8) (75- 1276), APC Anti-Human/Mouse CD11b (M1/70) (20-0112) (Tonbo); IL6 (11-7061-81) (Invitrogen); IL1β (31202) (Cell Signaling Technology); Brefeldin A Solution (420601), Monensin Solution (420701) (BioLegend); mouse TNF-α (430904) and IL-6 (431304) enzyme-linked immunosorbent assay (ELISA) kits were purchased from BioLegend; Griess Reagent System (G2930) was purchased from Promega.

### Cell culture

L929 cells were obtained from ATCC and maintained in DMEM (D5796) (Sigma) supplemented with 10% FBS (97068-085) (VWR). BMDMs were generated from bone marrow and cultured on Petri plates in DMEM containing 10% FBS plus 30% L929 culture media for 7 days. PerMacs were collected from peritoneal gavages 4 days after intraperitoneal injection of 3% brewer thioglycolate medium (B2551) (Sigma). Transfection of interferon-stimulatory DNA (ISD) (InvivoGen) into the cytosol of BMDMs was performed using Polyethyleneimine (PEI) (43896) (Alfa Aesar). Unless stated, 6 x 10^5^ BMDMs and 1.2 x10^6^ PerMacs per milliliter were used in in vitro experiments. The LPS-B5 Ultrapure (InvivoGen) concentration used was 200ng/ml for BMDMs and 20 ng/ml for PerMacs unless otherwise indicated. The 4-octyl Itaconate (Cayman) concentration was 125 μM and the Dimethyl fumarate (Sigma) concentration was 50 uM, and in experiments cells were exposed to both treatments 6 hrs before LPS stimulation.

### LPS in vivo challenge and multi-analyte cytokine analysis

Mice were intraperitoneally (i.p.) injected with LPS from Escherichia coli O55:B5 (L4005; Sigma). Blood was collected using EDTA-coated tubes at indicated time, then centrifuged at 1,000xg for 15 min at 4C°. The upper phase of plasma was saved and subjected to LEGENDplex Anti-Virus Response Panel (740446) (BioLegend) or LEGENDplex Mouse Inflammation Panel (740446) (BioLegend) for cytokine analysis.

### Blood and bone marrow staining for flow cytometry

Whole mouse blood was collected in sodium heparin tubes, and bone marrow was harvested from femurs and tibia. RBCs were lysed twice with ACK lysis buffer, and leukocytes were subjected to 1μg/ml LPS stimulation in the presence of brefeldin A and monensin for 4 hrs. Fc receptors were blocked with anti-Mouse CD16 / CD32 (2.4G2) antibody, cells stained with antibodies against surface proteins, permeabilized with Foxp3/Transcription Factor Staining Buffer Kit (TNB-0607-KIT) (Tonbo), and stained with antibodies against intracellular proteins. Cells were analyzed with a BD Fortessa X-20.

### *Listeria monocytogenes* infections

Age- and sex matched mice of indicated genotypes were used for infections, which were performed under BSL2 containment according to protocols approved by the Texas A&M IACUC. Two days prior to infection, cages’ plain water bottles were replaced with bottles containing 5 mg/ml streptomycin in water. The night before infection, chow was removed from the mouse cages. Mice were infected with log phase (O.D.600 = 0.5-1.0) *Listeria monocytogenes* (strain 10403S, gift from D. Portnoy) grown in BHI at 37°C. Bacteria were washed twice with warm, sterile PBS, and for each mouse, an inoculum of 1×10^8^ bacteria was placed on a 3 mm piece of bread with 3 ml of butter. Each mouse was individually fed one piece of Listeria-soaked bread and butter and then returned to their cage with fresh, antibiotic-free water and chow. Colonization was confirmed by assessing bacterial shedding in feces; at indicated time points, stools were collected from each mouse, dissolved in PBS, and spot plated as serial dilutions on LB plates.

Mice were weighed prior to infection and regularly throughout to monitor their health status. Bacterial burdens in spleens were determined by homogenizing spleens in 0.1% IGEPAL and plating serial dilutions on LB plates. Plasma was collected using EDTA-coated tubes at indicated time points for cytokine analyses.

### Quantitative PCR

To measure relative gene expression by qRT–PCR in cells and tissues, total cellular RNA was isolated using Quick-RNA microprep kit (Zymo Research). Approximately 0.5-1 ug RNA was isolated and cDNA was generated using the qScript cDNA Synthesis Kit (Quanta). cDNA was then subjected to qPCR using PerfeCTa SYBR Green FastMix (Quanta). Three technical replicates were performed for each biological sample, and expression values of each replicate were normalized against Gapdh or Rpl37 cDNA using the 2^-ΔΔCT^ method. For mtDNA abundance assessment, 2ng/μl of template DNA was used for qRT-PCR, and expression values of each replicate were normalized against nuclear-encoded Actb. All primer sequences used for qRT-PCR can be found in Extended Data Table 1.

### Immunoblotting

Cells and tissues were lysed in 1% NP-40 lysis buffer or 1% SDS lysis buffer supplemented with °C to obtain cellular lysate. After BCA protein assay (Thermo Fisher Scientific), equal amounts of protein (10–40□μg) were loaded into 10-20% SDS– PAGE gradient gels and transferred onto 0.22uM PVDF membranes. After air drying to return to a hydrophobic state, membranes were incubated in primary antibodies at 4°C overnight in 1X PBS containing 1% casein, HRP-conjugated secondary antibody at room temperature for 1 hour, and then developed with Luminata Crescendo Western HRP Substrate (Millipore).

### ELISA

Detached cells and debris were removed in cell culture supernatant or plasma prior to the assay by centrifugation. After incubating in capture antibodies at 4°C overnight, blocking with PBS containing 10% FBS at room temperature for 1 hr, standards and cell culture supernatant were added into the ELISA plates, followed by incubation of detection antibodies, Avidin-HPR and TMB for faster color development. Plates were washed with PBS containing 0.05% Tween-20 between each step.

### Immunofluorescence microscopy

Cells were grown on coverslips and treated as described. After washing in PBS, cells were fixed with 4% paraformaldehyde for 20 min, permeabilized with 0.1% Triton X-100 in PBS for 5 min, blocked with PBS containing 5% FBS for 30 min, stained with primary antibodies for 1h, and stained with secondary antibodies for 1h. Cells were washed with PBS containing 5% FBS between each step. Coverslips were mounted with ProLong Diamond Antifade Mountant with DAPI (Molecular Probes). Cells were imaged on a LSM 780 confocal microscope (Zeiss) with a 40 or 63x oil-immersed objective with Airyscan or a Cytation 5 (BioTek) with a 20x objective. 3-5 images per sample were acquired. At least 100 cells per genotype were used to obtain statistical significance for nuclear Nrf2 intensity analysis. Gen5 software (BioTek) was utilized to define nuclear region, by DAPI staining, and calculate the Nrf2 fluorescent intensity in the nuclear region of each cell.

### Metabolism assays

The Seahorse XFe96 Analyzer was used to measure mitochondrial respiration and glycolysis. Briefly, BMDMs and PerMacs were plated at a density of 5×10^4^ cells/well or 2×10^5^ cells/ well in 80µL of culture medium in Agilent Seahorse XF96 Cell Culture Microplate. DMF (50 uM) and/or LPS (10 ng/ml) were added 3 hrs after the cells were plated. Following a 16 hr incubation, cells were washed and replaced with XF assay medium (Base Medium Minimal DMEM supplemented with 2 mM Ala-Gln, pH 7.4) prior to analysis. Oxygen consumption rate (OCR) and extracellular acidification rate (ECAR) were measured after sequential addition of glucose 25 mM, oligomycin (1.5 uM), FCCP (1.5 µM)+ sodium pyruvate (1mM) and antimycin A (833 nM) and rotenone (833 nM). L-Lactate (700510), Aconitase (705502) (Cayman); LDH (MK401) (TaKaRa) assay kits were commercially purchased. Standard protocols provided with the kits were followed when performing all assays.

### Histology

Mice were euthanized and tissues were washed in PBS and incubated for 24L and then transferred to 70% ethanol. Tissue embedding, sectioning and staining were performed at AML Laboratories in Jacksonville, FL. Images of the H&E staining slides were acquired on Lionheart FX (BioTek) with a 20x or 40x objective. Images for myofiber width quantification were acquired on Nikon ECLIPSE Ts2 and a 40X objective. Ten cells from each sample, and three mice of each genotype were imaged and calculated blindly using NIS-Elements software (Nikon).

### Echocardiography

Mice were depilated at the chest area the day before echocardiogram measurement. Anesthesia was induced by placing mouse in a box with inhaled isoflurane at 2-3%. Afterwards, light anesthesia was maintained using a nose cone with inhaled isoflurane at 0.5-1%. Mice were immobilized on an instrumented and heated stage. Continuous electric cardiogram (ECG), respiration, and temperature (via rectal probe) were monitored. Light anesthesia and core temperature were maintained to ensure near physiological status with heart rate at range of 470-520 beats per minute. Transthoracic echocardiography was performed using a VisualSonics Vevo 3100 system with a MX550D imaging transducer (center frequency of 40 MHz). Parasternal long-axis (B-mode) and parasternal short-axis (M-mode) views of each animal were taken. Vevo LAB cardiac analysis package was used to analyze the data.

### RNA sequencing and bioinformatic analyses

Total cellular RNA from WT and *POLG* BMDMs and PerMacs was prepared using Quick-RNA microprep kit (Zymo Research) and used for the Next-generation RNA sequencing procedure at Texas A&M University Bioinformatics Core. RNA sequencing data were analyzed using BaseSpace Sequence Hub (Illmina). In brief, STAR algorithm of RNAseq Alignment V2.0.0 software was utilized to align the results to reference genome Mus musculus/mm10 (RefSeq), then RNA-seq Differential Expression V1.0.0 software was used to obtain raw gene expression files and determine statistically significant changes in gene expression in *POLG* macrophages relative to WT. Ingenuity Pathway Analysis software (QIAGEN) was used to identify gene families and putative upstream regulators in the datasets. Heat maps were generated using GraphPad Prism. Upon acceptance of the manuscript, datasets will be deposited in GEO according with NIH data sharing policies.

### Statistical analyses

Error bars displayed throughout the manuscript represent s.e.m. and were calculated from triplicate technical replicates of each biological sample unless otherwise indicated. For ex vivo experiments, error bars were calculated from the average of duplicate or triplicate technical replicates of at least 2 animals per point. For microscopy quantification, images were taken throughout the slide of each sample using DAPI channel to avoid bias of selection, and at least 100 cells per genotype were randomly selected to obtain statistical significance. To reduce potential experimental bias, samples for cardiac function (LVID;d, LVEF, IVS;s, LVID;s, Vol;s) and myofiber width analysis in Figure 6 were blinded to researchers when performing analysis. The identities were only revealed at the final data analysis stage. No randomization or blinding was used for all other animal studies. No statistical method was used to predetermine sample size. Data shown are representative of 2–3 independent experiments, including microscopy images, western blots, flow cytometry and all metabolism assays.

## Acknowledgments

**General**

We thank Dr. Rola Barhoumi Mouneimne for assistance with confocal microscopy, Ms. Robbie Moore for flow cytometry assistance, and Ms. Heidi Creed for assistance with the Vevo 3100 ultrasound.

## Author contributions

A.P.W and Y.L. designed the experiments, analyzed the data, and wrote the manuscript. Y.L. performed most of the experiments. C.G.M., S.T.O., C.E.B., and J.D.B. assisted with animal experiments, sample processing, and/or data analyses. S.L.B. and R.O.W. contributed to the *L. monocytogenes* infections, sample collection, and data analyses. C.W.T. and L.C.W. provided expertise and advice on cardiac measurements and flow cytometric experiments, respectively. A.P.W conceived the project and provided overall direction.

## Data and materials availability

All data needed to evaluate the conclusions drawn herein are present in the paper and/or the Supplementary Materials. Raw data will be provided upon request. RNA-Seq datasets will be deposited to GEO upon acceptance of the manuscript.

## Supplementary Materials

**Figure S1.**
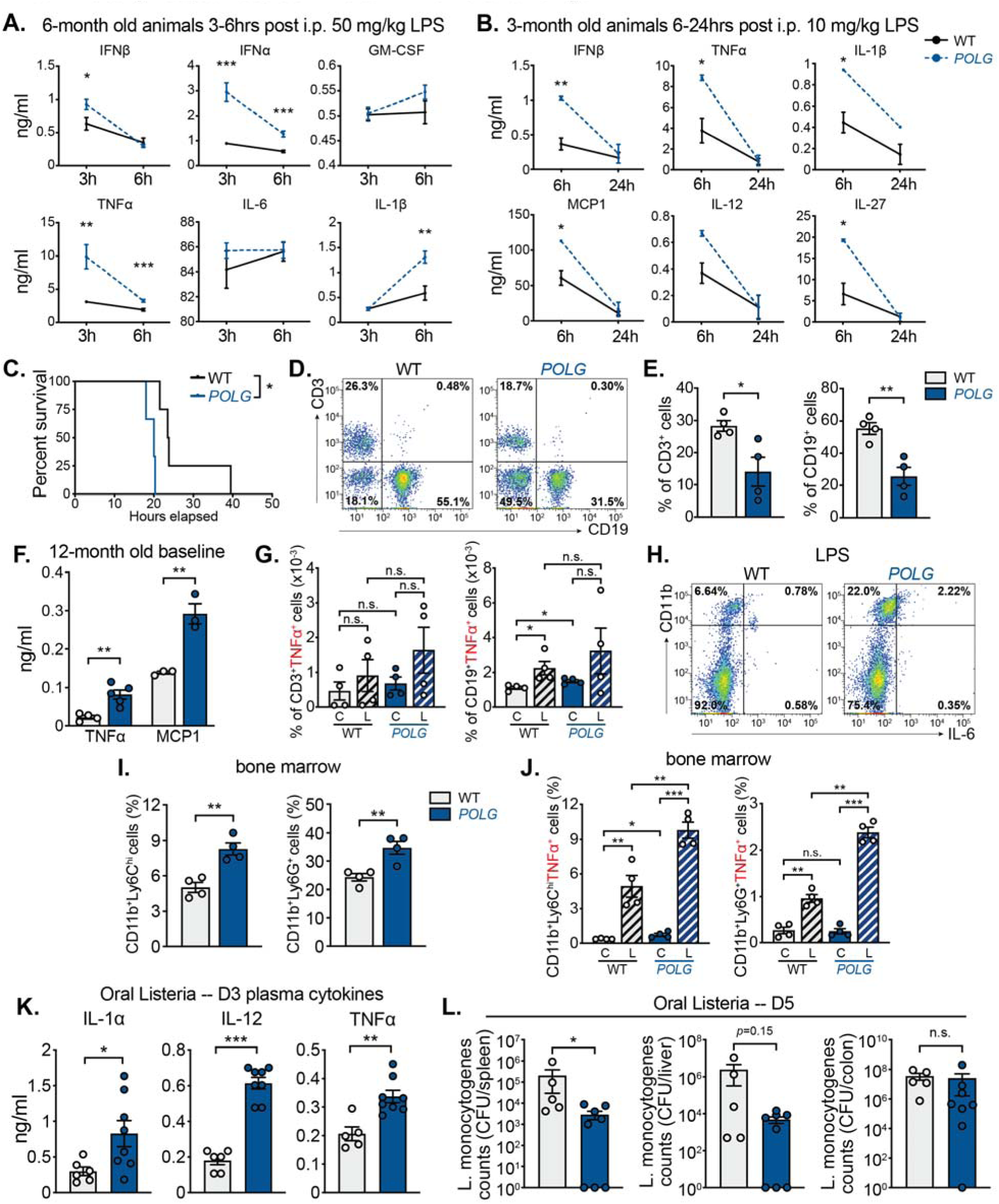
**CD11b^+^ myeloid cell expansion in POLG mutator mice drives systemic hyper-inflammatory responses to innate immune challenge.** (A and C) 6-month old WT and *POLG* mice were i.p. injected with 50 mg/kg LPS. Plasma was collected and subjected to multi-analyte cytokine analysis. (B) 3-month old WT and *POLG* mice were i.p. injected with 10 mg/kg LPS (n=3). Plasma was collected and subjected to multi-analyte cytokine analysis. (C) Kaplan-Meyer survival analysis was performed on 6-month old WT and *POLG* mice (n=3-4). Log-rank (Mantel-Cox) test was used to compare percent survival between different groups. (D and E) T cell (CD3^+^) and B cell (CD19^+^) populations in whole blood from 12-month old WT and *POLG* mice were evaluated by flow cytometry. Pseudocolor plots are representative of 4 independent experiments (D) and quantification is shown in (E). (F) Cytokine levels in WT and *POLG* mutator plasma at baseline. (G) CD3^+^TNFα^+^ and CD19^+^TNFα^+^ MFI in unstimulated [C] or LPS challenged [L] whole blood from 12-month old WT and *POLG* mice as determined by flow cytometry. (H) CD11b^+^IL-6^+^ population in LPS challenged whole blood from 12-month old WT and *POLG* mice was evaluated by flow cytometry. Pseudocolor plots are representative of 4 independent experiments. (I) The percentage of CD11b^+^Ly6C^hi^ and CD11b^+^Ly6G^+^ cells in WT and *POLG* bone marrow as determined by flow cytometry. (J) The percentage of CD11b^+^Ly6C^hi^TNFα^+^ and CD11b^+^Ly6G^+^TNFα^+^ populations in unstimulated [C] or LPS challenged [L] bone marrow from 12-month old WT and *POLG* mice as quantified by flow cytometry. (K) Cytokine levels in WT and *POLG* mutator plasma 3 days post *L. monocytogenes* infection. (L) *L. monocytogenes* CFU counts in mouse spleen, liver and colon D5 post infection. Mann-Whitney test was used to compare CFUs in tissues between WT and POLG mice. Unless stated, statistical significance was determined using unpaired Student’s t-tests. \**P* < 0.05; \*\**P* < 0.01; \*\*\**P* < 0.001. Error bars represent S.E.M.

**Figure S2.**
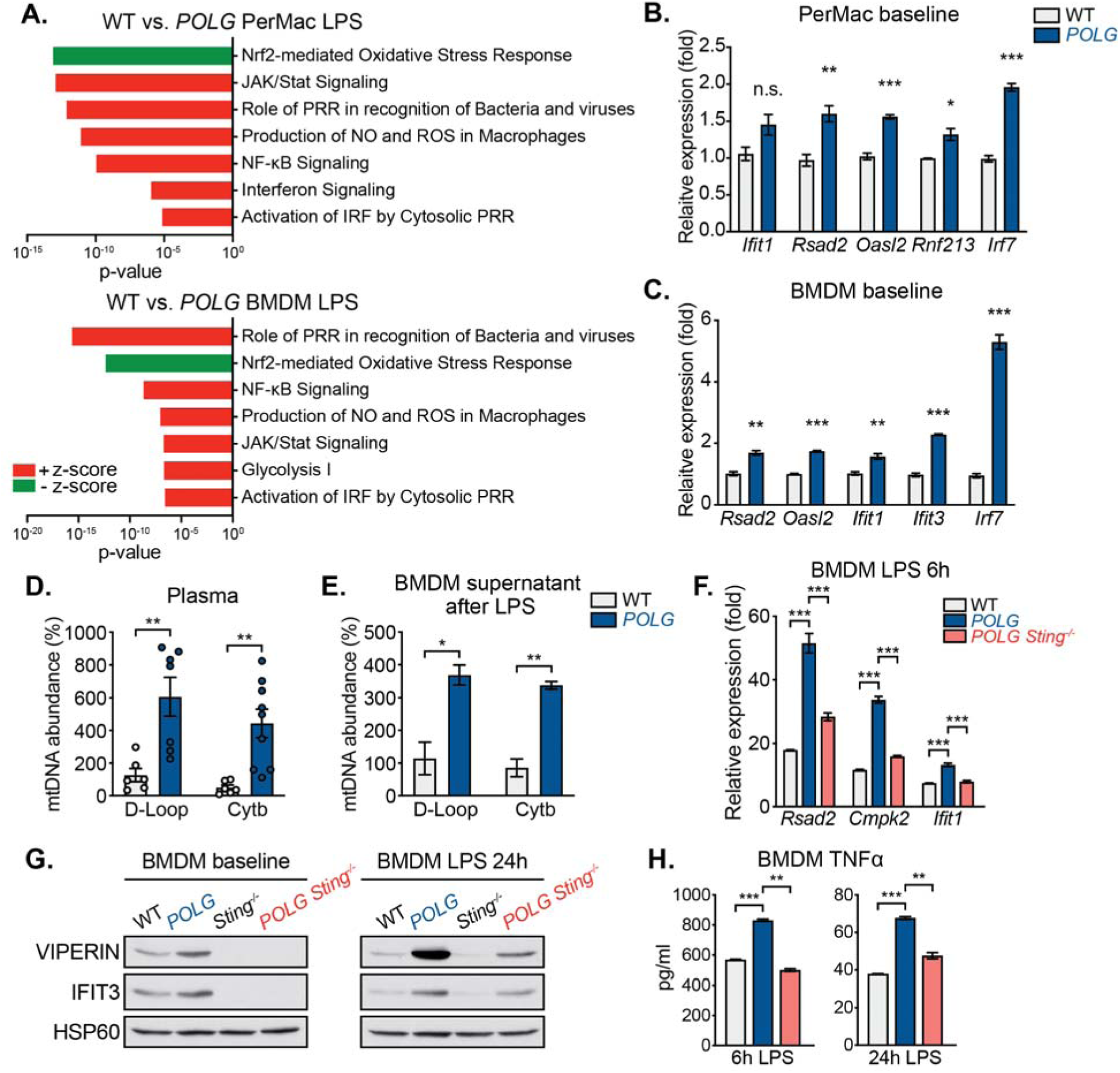
**STING regulates enhanced IFN-I and pro-inflammatory responses in LPS challenged POLG mutator macrophages.** (A) Ingenuity Pathway Analysis comparing RNAseq datasets from LPS treated (200ng/ml for 6h) WT and *POLG* PerMacs and BMDMs. (B and C) qRT-PCR analysis of ISG expression in resting WT and *POLG* PerMacs (B) and BMDMs (C). (D) qPCR analysis of circulating, cell-free mtDNA abundance in 12-month old WT and *POLG* mutator plasma. n=6-9 animals per genotype. (E) qPCR analysis of WT and *POLG* BMDMs mtDNA release into the culture supernatant 48h after 200ng/ml LPS exposure. (F) qRT-PCR analysis of ISG expression in WT, *POLG* and *POLG Sting^-/-^* BMDMs after LPS challenge. (G) Protein abundance of ISGs in WT, *POLG*, *Sting^-/-^* and *POLG Sting^-/-^* BMDMs without or with LPS treatment. (H) Pro-inflammatory cytokine secretion of WT, *POLG* and *POLG Sting^-/-^* BMDMs after LPS challenge. Statistical significance was determined using unpaired Student’s t-tests. \**P* < 0.05; \*\**P* < 0.01; \*\*\**P* < 0.001. Error bars represent S.E.M.

**Figure S3.**
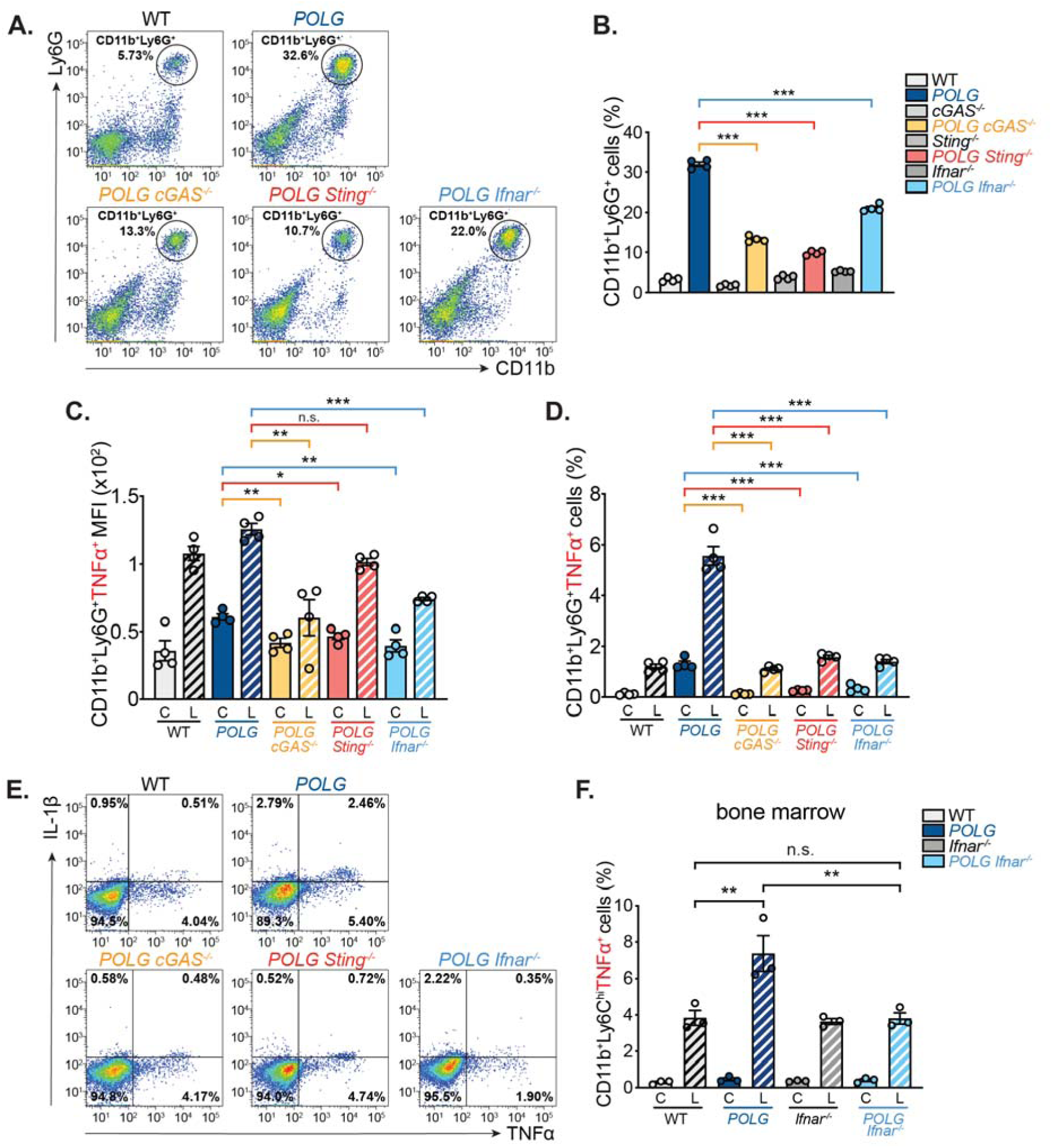
**Neutrophil expansion and elevated pro-inflammatory cytokine production in POLG mutator mice is regulated by the cGAS-STING-IFN-I signaling axis.** (A and B) CD11b^+^Ly6G^+^ neutrophil population in the whole blood was evaluated by flow cytometry. Pseudocolor plots are representative of 4 independent experiments (A) and quantification of the percentage of CD11b^+^Ly6G^+^ cells is shown in (B). (C and D) CD11b^+^Ly6G^+^TNFα^+^ neutrophil population in unstimulated [C] or LPS challenged [L] whole blood from 12-month old cohorts was evaluated by flow cytometry. Quantification of CD11b^+^Ly6G^+^TNFα^+^ mean fluorescent intensity (MFI) is shown in (C), and the percentage of CD11b^+^Ly6G^+^TNFα^+^ cells is shown in (D). (E) Intracellular TNFα and IL-1β staining was performed on LPS challenged whole blood from 12-month old cohorts. Pseudocolor plots are representative of 2 independent experiments. (F) The percentage of CD11b^+^Ly6C^hi^TNFα^+^ inflammatory monocyte population in unstimulated [C] or LPS challenged [L] bone marrow from 12-month old cohorts as quantified by flow cytometry. Statistical significance was determined using unpaired Student’s t-tests or ANOVA. \**P* < 0.05; \*\**P* < 0.01; \*\*\**P* < 0.001. Error bars represent S.E.M.

**Figure S4.**
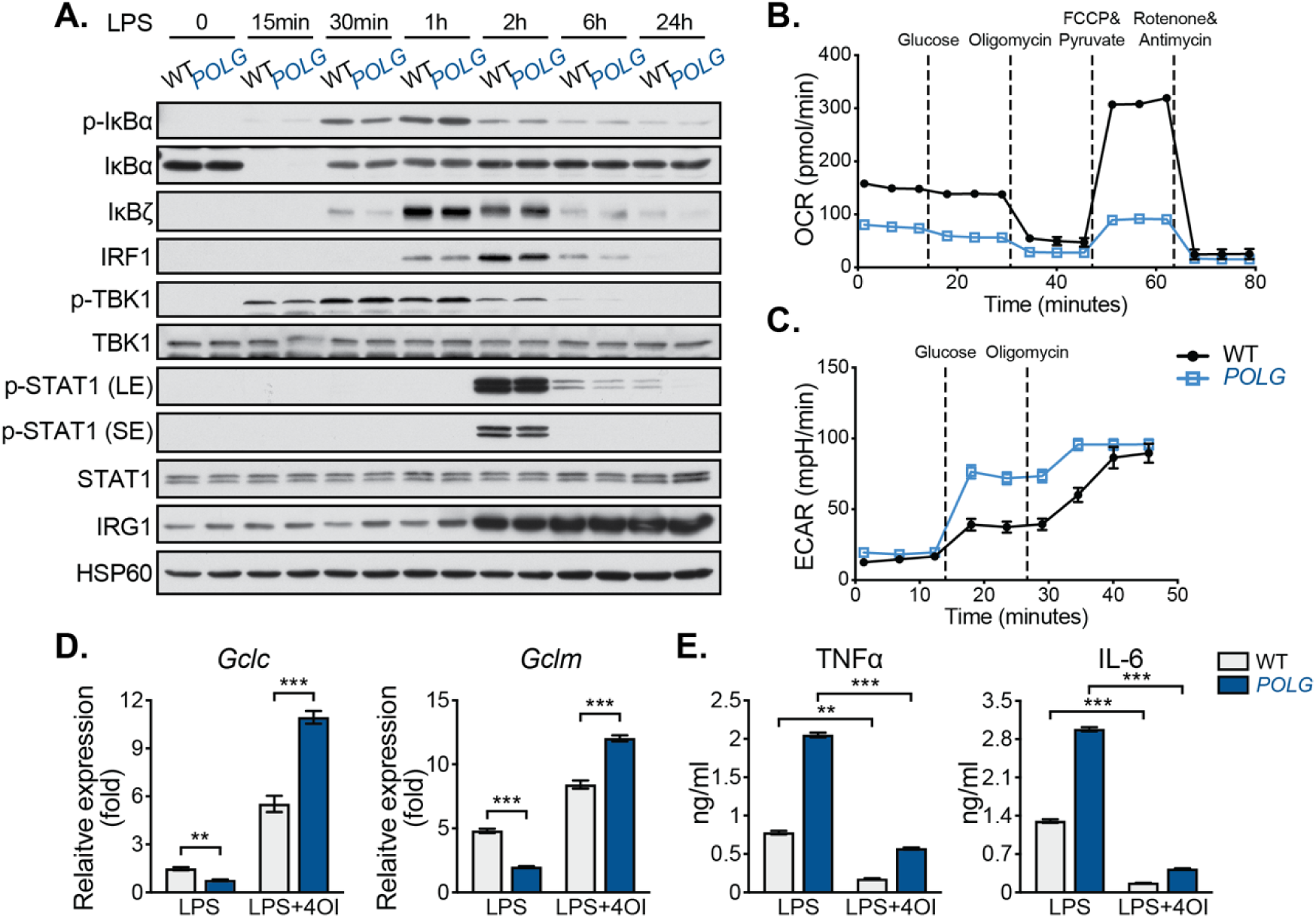
**POLG mutator macrophages exhibit normal NF-κB and IFN-I signaling kinetics in response to LPS, but displayed impaired Nrf2 activity.** (A) NF-κB and IFN-I signaling pathways protein expression in WT and *POLG* BMDMs after LPS exposure over a time-course. (B and C) Seahorse analysis of resting WT and *POLG* BMDMs represented as OCR (B) and ECAR (C). (D) qRT-PCR analysis of Nrf2 target gene expression in WT and *POLG* PerMacs after LPS or LPS+4OI treatment. (E) Pro-inflammatory cytokine secretion by WT and *POLG* PerMacs after LPS or LPS+4OI treatment. Statistical significance was determined using unpaired Student’s t-tests. \**P* < 0.05; \*\**P* < 0.01; \*\*\**P* < 0.001. Error bars represent S.E.M.

**Figure S5.**
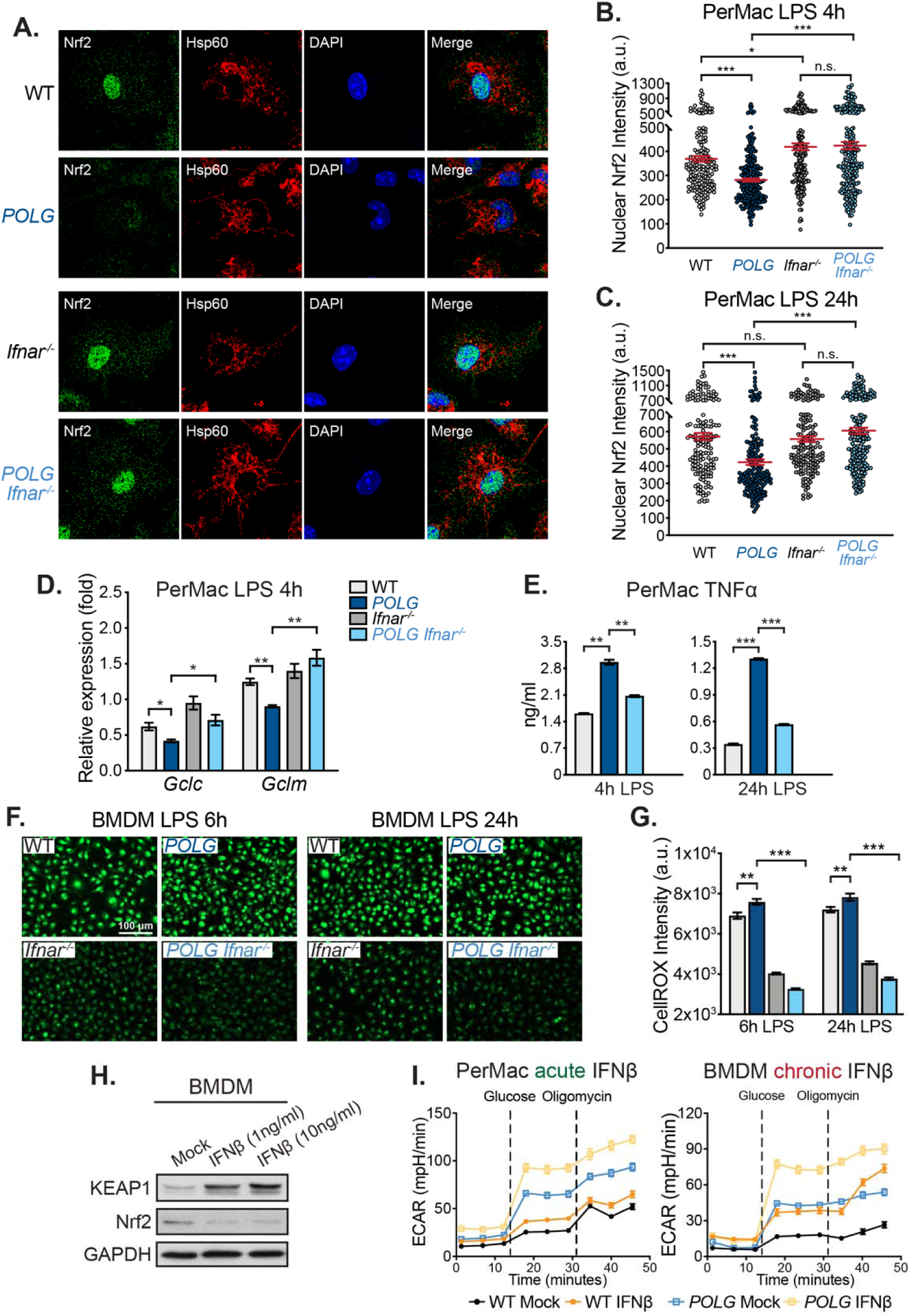
**Elevated IFN-I signaling represses Nrf2 activity and drives pro-inflammatory cytokine and metabolic phenotypes in POLG mutator macrophages.** (A) Representative confocal microscopy images of LPS treated PerMacs stained with anti-Nrf2 and -Hsp60 antibodies and DAPI. (B and C) Quantification of nuclear Nrf2 staining intensity in PerMacs 4h (b) or 24h (c) after LPS exposure. a.u., arbitrary unit. (D) Nrf2 target gene expression in PerMacs post LPS exposure. (E) TNFα secretion by PerMacs 4h and 24h post LPS exposure. (F) Representative microscopy images of LPS treated BMDMs stained with CellROX. (G) Quantification of MFI of CellROX staining in LPS treated BMDMs. (H) Protein expression in WT BMDM after 24h of IFNβ exposure. (I) Seahorse ECAR of WT and *POLG* PerMacs and BMDMs post-acute (24h) or -chronic (5 days) IFNβ exposure. Statistical significance was determined using unpaired Student’s t-tests or ANOVA. \**P* < 0.05; \*\**P* < 0.01; \*\*\**P* < 0.001. Error bars represent S.E.M.

**Figure S6.**
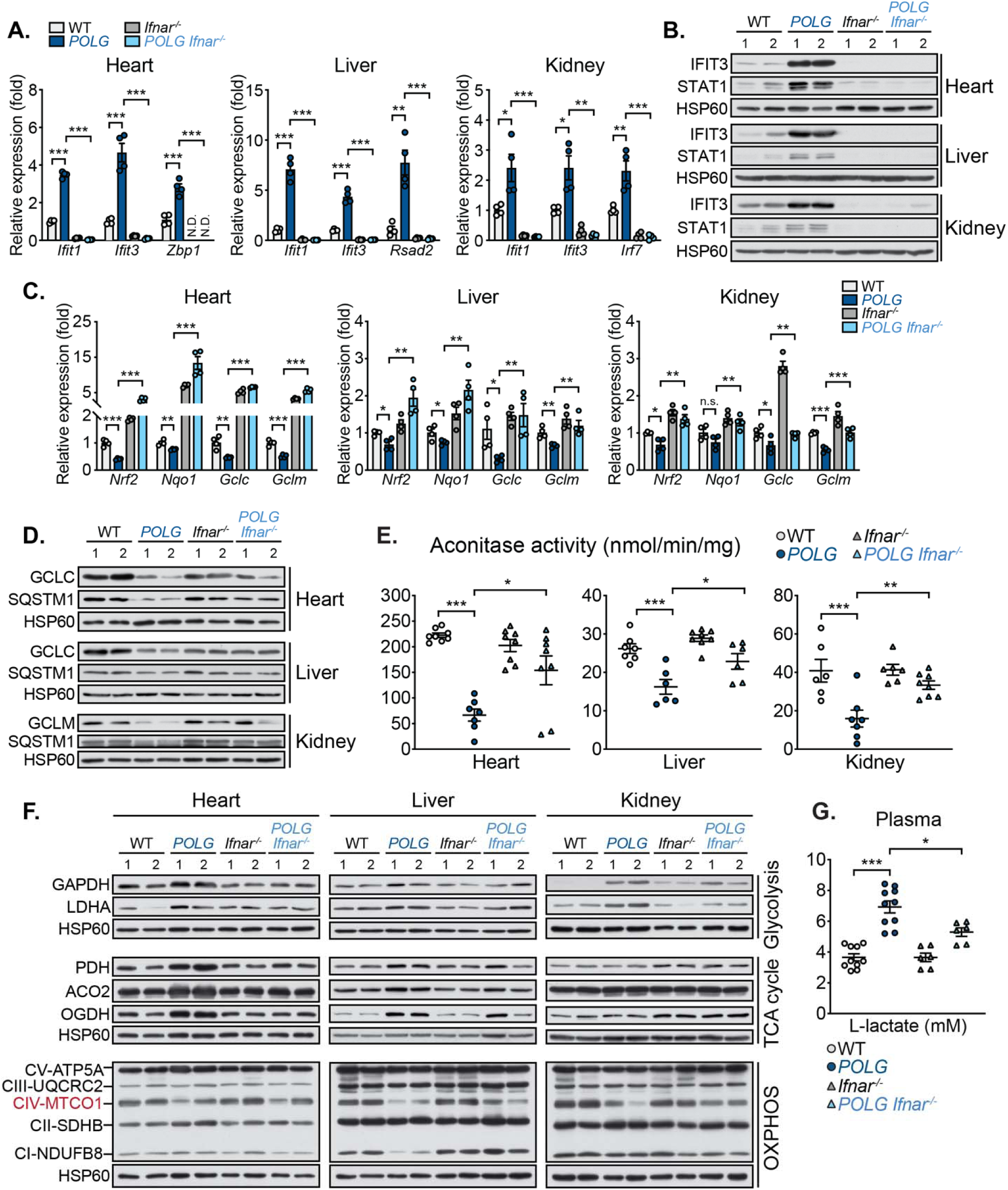
**Chronic IFN-I signaling in aged POLG mutator mice represses Nrf2 target gene expression and elevates oxidative stress in tissues.** (A and B) Transcript (A) and protein (B) levels of ISGs in WT, *POLG*, *Ifnar^-/-^* and *POLG Ifnar^-/-^* mouse heart, liver and kidney. 2 animals (12-month old) per genotype are represented. (C and D) Transcript (C) and protein (D) levels of and Nrf2-targets in WT, *POLG*, *Ifnar^-/-^* and *POLG Ifnar^-/-^* mouse heart, liver and kidney. 2 animals (12-month old) per genotype are represented. (E) Aconitase activity in WT, *POLG*, *Ifnar^-/-^* and *POLG Ifnar^-/-^* heart, liver and kidney. n=4-5 animals per genotype and each animal was represented in duplicates. (F) Protein levels of glycolytic enzymes, TCA cycle enzymes, and OXPHOS complexes in WT, *POLG*, *Ifnar^-/-^* and *POLG Ifnar^-/-^* mouse tissues. 2 animals (12-month old) per genotype are represented. (G) L-lactate concentration in plasma from WT, *POLG*, *Ifnar^-/-^* and *POLG Ifnar^-/-^* mice. n=6-10 animals per genotype. Statistical significance was determined using unpaired Student’s t-tests. \**P* < 0.05; \*\**P* < 0.01; \*\*\**P* < 0.001. Error bars represent S.E.M.

**Figure S7.**
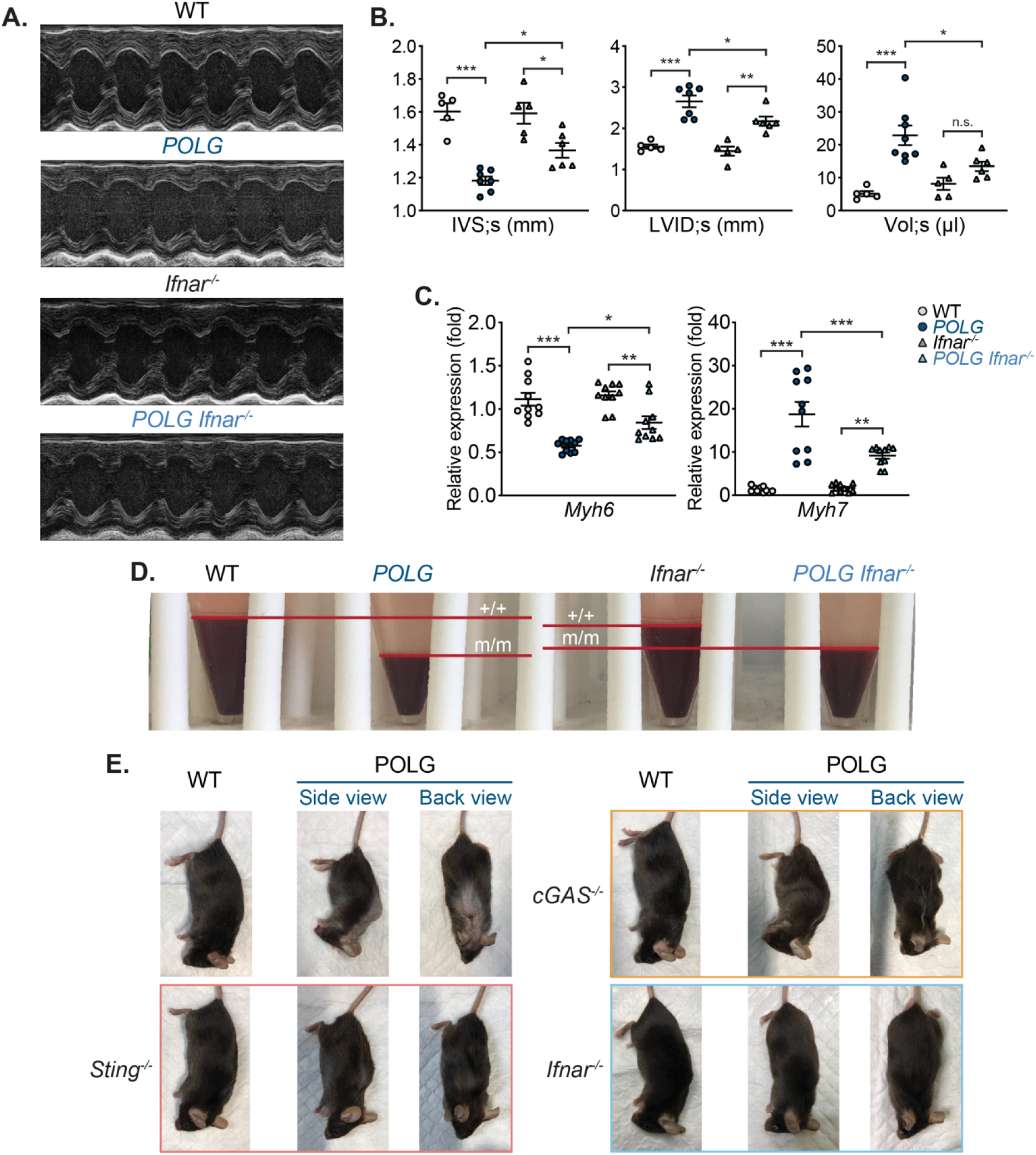
**Ablation of IFN-I signaling attenuates cardiomyopathy and anemia, and improves body condition of POLG mutators.** (A) Representative M-mode ultrasound echocardiograms of WT, *POLG*, *Ifnar^-/-^* and *POLG Ifnar^- /-^* hearts. (B) Interventricular septum thickness at end-systole (IVS;s), left ventricular internal dimension at end-systole (LVID;s) and left ventricular volume at end-systole (Vol;s) were measured in WT, *POLG*, *Ifnar^-/-^* and *POLG Ifnar^-/-^* mouse hearts. n=5-8 animals per genotype. (C) qRT-PCR analysis of *Myh6* and *Myh7* expression from heart homogenates. n=5 animals per genotype and each animal was represented in duplicates. (D) Representative images of blood pellets from exsanguinated 12-month old mice. (E) Representative images of 12-month old *POLG*, *POLG cGAS^-/-^*, *POLG Sting^-/-^* and *POLG Ifnar^-/-^* animals. Statistical significance was determined using ANOVA.\**P* < 0.05; \*\**P* < 0.01; \*\*\**P* < 0.001. Error bars represent S.E.M.

**Figure S8.**
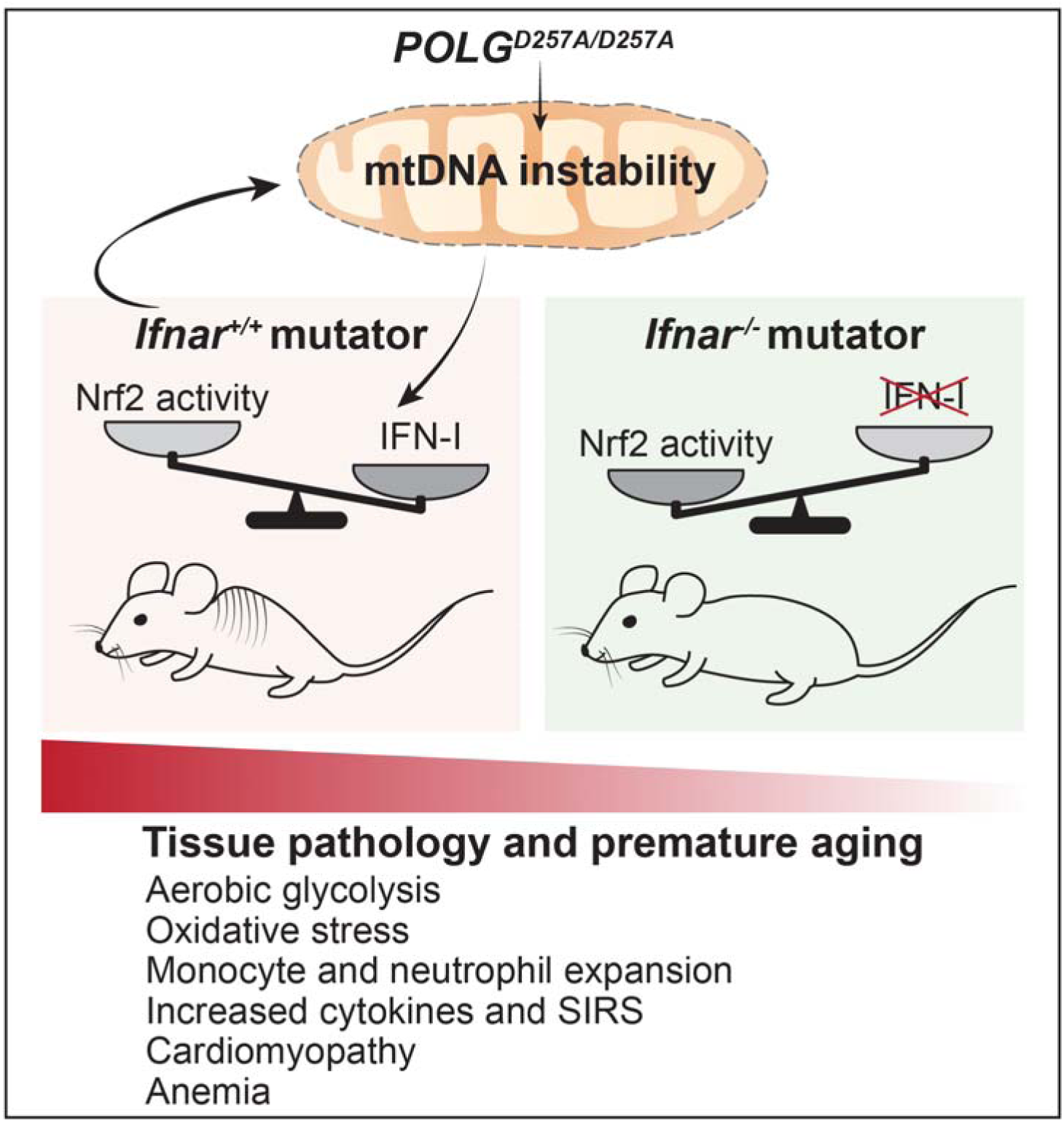
**Imbalances in IFN-I and Nrf2 signaling contribute to inflammatory and age- related phenotypes in POLG mutator mice.** mtDNA instability and mitochondrial dysfunction in POLG mutator mice lead to elevated IFN-I responses, which subsequently repress Nrf2 activity and enhance aerobic glycolysis. Consequently, chronic IFN-I responses augment the expansion and inflammatory potential of CD11b^+^ myeloid cells and macrophages, while also contributing to cardiomyopathy and anemia in mutator mice. Genetic ablation of IFN-I signaling relieves the break on Nrf2 and significantly improves healthspan by limiting myeloid reprogramming, inflammation, and tissue dysfunction in mtDNA mutator mice.

## References and Notes

1. A. P. West, Mitochondrial dysfunction as a trigger of innate immune responses and inflammation. Toxicology. 391, 54–63 (2017).

2. A. P. West, G. S. Shadel, S. Ghosh, Mitochondria in innate immune responses. Nat Rev Immunol. 11, 389–402 (2011).

3. S. E. Weinberg, L. A. Sena, N. S. Chandel, Mitochondria in the regulation of innate and adaptive immunity. Immunity. 42, 406–417 (2015).

4. E. L. Mills, B. Kelly, L. A. J. O’Neill, Mitochondria are the powerhouses of immunity. Nature Immunology. 18, 488–498 (2017).

5. R. J. Youle, Mitochondria-Striking a balance between host and endosymbiont. Science. 365 (2019), doi:10.1126/science.aaw9855.

6. A. P. West, G. S. Shadel, Mitochondrial DNA in innate immune responses and inflammatory pathology. Nat Rev Immunol. 17, 363–375 (2017).

7. A. P. West, W. Khoury-Hanold, M. Staron, M. C. Tal, C. M. Pineda, S. M. Lang, M. Bestwick, B. A. Duguay, N. Raimundo, D. A. MacDuff, S. M. Kaech, J. R. Smiley, R. E. Means, A. Iwasaki, G. S. Shadel, Mitochondrial DNA stress primes the antiviral innate immune response. Nature. 520, 553–557 (2015).

8. A. Ablasser, Z. J. Chen, cGAS in action: Expanding roles in immunity and inflammation. Science. 363 (2019), doi:10.1126/science.aat8657.

9. K. Nakahira, S. Hisata, A. M. K. Choi, The Roles of Mitochondrial Damage-Associated Molecular Patterns in Diseases. Antioxid. Redox Signal. 23, 1329–1350 (2015).

10. Z. Wu, S. Oeck, A. P. West, K. C. Mangalhara, A. G. Sainz, L. E. Newman, X.-O. Zhang, L. Wu, Q. Yan, M. Bosenberg, Y. Liu, P. L. Sulkowski, V. Tripple, S. M. Kaech, P. M. Glazer, G. S. Shadel, Mitochondrial DNA stress signalling protects the nuclear genome. Nature Metabolism. 1, 1209–1218 (2019).

11. G. S. Gorman, P. F. Chinnery, S. DiMauro, M. Hirano, Y. Koga, R. McFarland, A. Suomalainen, D. R. Thorburn, M. Zeviani, D. M. Turnbull, Mitochondrial diseases. Nature Reviews Disease Primers. 2, 1–22 (2016).

12. J. D. Stumpf, R. P. Saneto, W. C. Copeland, Clinical and Molecular Features of POLG-Related Mitochondrial Disease. Cold Spring Harb Perspect Biol. 5 (2013), doi:10.1101/cshperspect.a011395.

13. S. Eom, H. N. Lee, S. Lee, H.-C. Kang, J. S. Lee, H. D. Kim, Y.-M. Lee, Cause of Death in Children With Mitochondrial Diseases. Pediatr. Neurol. 66, 82–88 (2017).

14. S. M. Kapnick, S. E. Pacheco, P. J. McGuire, The emerging role of immune dysfunction in mitochondrial diseases as a paradigm for understanding immunometabolism. Metab. Clin. Exp. 81, 97–112 (2018).

15. M. A. Walker, N. Slate, A. Alejos, S. Volpi, R. S. Iyengar, D. Sweetser, K. B. Sims, J. E. Walter, Predisposition to infection and SIRS in mitochondrial disorders: 8 years’ experience in an academic center. Journal of Allergy and Clinical Immunology. In Practice; Amsterdam. 2, 465–468 (2014).

16. T. N. Tarasenko, S. E. Pacheco, M. K. Koenig, J. Gomez-Rodriguez, S. M. Kapnick, F. Diaz, P. M. Zerfas, E. Barca, J. Sudderth, R. J. DeBerardinis, R. Covian, R. S. Balaban, S. DiMauro, P. J. McGuire, Cytochrome c Oxidase Activity Is a Metabolic Checkpoint that Regulates Cell Fate Decisions During T Cell Activation and Differentiation. Cell Metab. 25, 1254–1268.e7 (2017).

17. O. Hasselmann, N. Blau, V. T. Ramaekers, E. V. Quadros, J. M. Sequeira, M. Weissert, Cerebral folate deficiency and CNS inflammatory markers in Alpers disease. Mol. Genet. Metab. 99, 58–61 (2010).

18. H. M. Wilkins, I. W. Weidling, Y. Ji, R. H. Swerdlow, Mitochondria-Derived Damage-Associated Molecular Patterns in Neurodegeneration. Front Immunol. 8, 508 (2017).

19. M. J. Young, W. C. Copeland, Human mitochondrial DNA replication machinery and disease. Curr Opin Genet Dev. 38, 52–62 (2016).

20. S. Rahman, W. C. Copeland, POLG-related disorders and their neurological manifestations. Nat Rev Neurol. 15, 40–52 (2019).

21. B. Singh, K. M. Owens, P. Bajpai, M. M. Desouki, V. Srinivasasainagendra, H. K. Tiwari, K. K. Singh, Mitochondrial DNA Polymerase POLG1 Disease Mutations and Germline Variants Promote Tumorigenic Properties. PLoS ONE. 10, e0139846 (2015).

22. G. Davidzon, P. Greene, M. Mancuso, K. J. Klos, J. E. Ahlskog, M. Hirano, S. DiMauro, Early-onset familial parkinsonism due to POLG mutations. Ann. Neurol. 59, 859–862 (2006).

23. G. C. Kujoth, A. Hiona, T. D. Pugh, S. Someya, K. Panzer, S. E. Wohlgemuth, T. Hofer, A. Y. Seo, R. Sullivan, W. A. Jobling, J. D. Morrow, H. V. Remmen, J. M. Sedivy, T. Yamasoba, M. Tanokura, R. Weindruch, C. Leeuwenburgh, T. A. Prolla, Mitochondrial DNA Mutations, Oxidative Stress, and Apoptosis in Mammalian Aging. Science. 309, 481– 484 (2005).

24. A. Trifunovic, A. Wredenberg, M. Falkenberg, J. N. Spelbrink, A. T. Rovio, C. E. Bruder, M. Bohlooly-Y, S. Gidlöf, A. Oldfors, R. Wibom, J. Törnell, H. T. Jacobs, N.-G. Larsson, Premature ageing in mice expressing defective mitochondrial DNA polymerase. Nature. 429, 417–423 (2004).

25. K. Szczepanowska, A. Trifunovic, Different faces of mitochondrial DNA mutators. Biochimica et Biophysica Acta (BBA) - Bioenergetics. 1847, 1362–1372 (2015).

26. J. Y. Jang, A. Blum, J. Liu, T. Finkel, The role of mitochondria in aging. J. Clin. Invest. 128, 3662–3670 (2018).

27. A. Logan, I. G. Shabalina, T. A. Prime, S. Rogatti, A. V. Kalinovich, R. C. Hartley, R. C. Budd, B. Cannon, M. P. Murphy, In vivo levels of mitochondrial hydrogen peroxide increase with age in mtDNA mutator mice. Aging Cell. 13, 765–768 (2014).

28. M. L. Chen, T. D. Logan, M. L. Hochberg, S. G. Shelat, X. Yu, G. E. Wilding, W. Tan, G. C. Kujoth, T. A. Prolla, M. A. Selak, M. Kundu, M. Carroll, J. E. Thompson, Erythroid dysplasia, megaloblastic anemia, and impaired lymphopoiesis arising from mitochondrial dysfunction. Blood. 114, 4045–4053 (2009).

29. A. Safdar, J. M. Bourgeois, D. I. Ogborn, J. P. Little, B. P. Hettinga, M. Akhtar, J. E. Thompson, S. Melov, N. J. Mocellin, G. C. Kujoth, T. A. Prolla, M. A. Tarnopolsky, Endurance exercise rescues progeroid aging and induces systemic mitochondrial rejuvenation in mtDNA mutator mice. Proc. Natl. Acad. Sci. U.S.A. 108, 4135–4140 (2011).

30. C. Shi, T. M. Hohl, I. Leiner, M. J. Equinda, X. Fan, E. G. Pamer, Ly6G+ neutrophils are dispensable for defense against systemic Listeria monocytogenes infection. J Immunol. 187, 5293–5298 (2011).

31. H. Maekawa, T. Inoue, H. Ouchi, T.-M. Jao, R. Inoue, H. Nishi, R. Fujii, F. Ishidate, T. Tanaka, Y. Tanaka, N. Hirokawa, M. Nangaku, R. Inagi, Mitochondrial Damage Causes Inflammation via cGAS-STING Signaling in Acute Kidney Injury. Cell Rep. 29, 1261–1273.e6 (2019).

32. K. W. Chung, P. Dhillon, S. Huang, X. Sheng, R. Shrestha, C. Qiu, B. A. Kaufman, J. Park, L. Pei, J. Baur, M. Palmer, K. Susztak, Mitochondrial Damage and Activation of the STING Pathway Lead to Renal Inflammation and Fibrosis. Cell Metabolism. 30, 784–799.e5 (2019).

33. D. A. Sliter, J. Martinez, L. Hao, X. Chen, N. Sun, T. D. Fischer, J. L. Burman, Y. Li, Z. Zhang, D. P. Narendra, H. Cai, M. Borsche, C. Klein, R. J. Youle, Parkin and PINK1 mitigate STING-induced inflammation. Nature. 561, 258–262 (2018).

34. G. Rackov, R. Shokri, M. Á. De Mon, C. Martínez-A., D. Balomenos, The Role of IFN-β during the Course of Sepsis Progression and Its Therapeutic Potential. Front Immunol. 8 (2017), doi:10.3389/fimmu.2017.00493.

35. S. H. Park, K. Kang, E. Giannopoulou, Y. Qiao, K. Kang, G. Kim, K.-H. Park-Min, L. B. Ivashkiv, Type I interferons and the cytokine TNF cooperatively reprogram the macrophage epigenome to promote inflammatory activation. Nature Immunology. 18, 1104–1116 (2017).

36. M. Karaghiosoff, R. Steinborn, P. Kovarik, G. Kriegshäuser, M. Baccarini, B. Donabauer, U. Reichart, T. Kolbe, C. Bogdan, T. Leanderson, D. Levy, T. Decker, M. Müller, Central role for type I interferons and Tyk2 in lipopolysaccharide-induced endotoxin shock. Nature Immunology. 4, 471–477 (2003).

37. M. B. Buechler, T. H. Teal, K. B. Elkon, J. A. Hamerman, Cutting Edge: Type I IFN Drives Emergency Myelopoiesis and Peripheral Myeloid Expansion during Chronic TLR7 Signaling. The Journal of Immunology. 190, 886–891 (2013).

38. P. Y. Lee, Y. Li, Y. Kumagai, Y. Xu, J. S. Weinstein, E. S. Kellner, D. C. Nacionales, E. J. Butfiloski, N. van Rooijen, S. Akira, E. S. Sobel, M. Satoh, W. H. Reeves, Type I interferon modulates monocyte recruitment and maturation in chronic inflammation. Am. J. Pathol. 175, 2023–2033 (2009).

39. S.-U. Seo, H.-J. Kwon, H.-J. Ko, Y.-H. Byun, B. L. Seong, S. Uematsu, S. Akira, M.-N. Kweon, Type I Interferon Signaling Regulates Ly6Chi Monocytes and Neutrophils during Acute Viral Pneumonia in Mice. PLOS Pathogens. 7, e1001304 (2011).

40. R. Channappanavar, A. R. Fehr, R. Vijay, M. Mack, J. Zhao, D. K. Meyerholz, S. Perlman, Dysregulated Type I Interferon and Inflammatory Monocyte-Macrophage Responses Cause Lethal Pneumonia in SARS-CoV-Infected Mice. Cell Host Microbe. 19, 181–193 (2016).

41. O. Majer, C. Bourgeois, F. Zwolanek, C. Lassnig, D. Kerjaschki, M. Mack, M. Müller, K. Kuchler, Type I Interferons Promote Fatal Immunopathology by Regulating Inflammatory Monocytes and Neutrophils during Candida Infections. PLOS Pathogens. 8, e1002811 (2012).

42. E. H. Kobayashi, T. Suzuki, R. Funayama, T. Nagashima, M. Hayashi, H. Sekine, N. Tanaka, T. Moriguchi, H. Motohashi, K. Nakayama, M. Yamamoto, Nrf2 suppresses macrophage inflammatory response by blocking proinflammatory cytokine transcription. Nature Communications. 7, 1–14 (2016).

43. E. L. Mills, D. G. Ryan, H. A. Prag, D. Dikovskaya, D. Menon, Z. Zaslona, M. P. Jedrychowski, A. S. H. Costa, M. Higgins, E. Hams, J. Szpyt, M. C. Runtsch, M. S. King, J. F. McGouran, R. Fischer, B. M. Kessler, A. F. McGettrick, M. M. Hughes, R. G. Carroll, L. M. Booty, E. V. Knatko, P. J. Meakin, M. L. J. Ashford, L. K. Modis, G. Brunori, D. C. Sévin, P. G. Fallon, S. T. Caldwell, E. R. S. Kunji, E. T. Chouchani, C. Frezza, A. T. Dinkova-Kostova, R. C. Hartley, M. P. Murphy, L. A. O’Neill, Itaconate is an anti-inflammatory metabolite that activates Nrf2 via alkylation of KEAP1. Nature. 556, 113–117 (2018).

44. K. Taguchi, H. Motohashi, M. Yamamoto, Molecular mechanisms of the Keap1–Nrf2 pathway in stress response and cancer evolution. Genes Cells. 16, 123–140 (2011).

45. M. D. Kornberg, P. Bhargava, P. M. Kim, V. Putluri, A. M. Snowman, N. Putluri, P. A. Calabresi, S. H. Snyder, Dimethyl fumarate targets GAPDH and aerobic glycolysis to modulate immunity. Science. 360, 449–453 (2018).

46. D. Olagnier, R. R. Lababidi, S. B. Hadj, A. Sze, Y. Liu, S. D. Naidu, M. Ferrari, Y. Jiang, C. Chiang, V. Beljanski, M.-L. Goulet, E. V. Knatko, A. T. Dinkova-Kostova, J. Hiscott, R. Lin, Activation of Nrf2 Signaling Augments Vesicular Stomatitis Virus Oncolysis via Autophagy-Driven Suppression of Antiviral Immunity. Mol. Ther. 25, 1900–1916 (2017).

47. C. Gunderstofte, M. B. Iversen, S. Peri, A. Thielke, S. Balachandran, C. K. Holm, D. Olagnier, Nrf2 Negatively Regulates Type I Interferon Responses and Increases Susceptibility to Herpes Genital Infection in Mice. Front Immunol. 10, 2101 (2019).

48. D. Olagnier, A. M. Brandtoft, C. Gunderstofte, N. L. Villadsen, C. Krapp, A. L. Thielke, A. Laustsen, S. Peri, A. L. Hansen, L. Bonefeld, J. Thyrsted, V. Bruun, M. B. Iversen, L. Lin, V. M. Artegoitia, C. Su, L. Yang, R. Lin, S. Balachandran, Y. Luo, M. Nyegaard, B. Marrero, R. Goldbach-Mansky, M. Motwani, D. G. Ryan, K. A. Fitzgerald, L. A. O’Neill, A. K. Hollensen, C. K. Damgaard, F. v de Paoli, H. C. Bertram, M. R. Jakobsen, T. B. Poulsen, C. K. Holm, Nrf2 negatively regulates STING indicating a link between antiviral sensing and metabolic reprogramming. Nature Communications. 9, 1–13 (2018).

49. N. J. Hos, R. Ganesan, S. Gutiérrez, D. Hos, J. Klimek, Z. Abdullah, M. Krönke, N. Robinson, Type I interferon enhances necroptosis of Salmonella Typhimurium-infected macrophages by impairing antioxidative stress responses. J. Cell Biol. 216, 4107–4121 (2017).

50. M. Riedelberger, P. Penninger, M. Tscherner, M. Seifert, S. Jenull, C. Brunnhofer, B. Scheidl, I. Tsymala, C. Bourgeois, A. Petryshyn, W. Glaser, A. Limbeck, B. Strobl, G. Weiss, K. Kuchler, Type I Interferon Response Dysregulates Host Iron Homeostasis and Enhances Candida glabrata Infection. Cell Host & Microbe. 27, 454–466.e8 (2020).

51. S. Kovac, P. R. Angelova, K. M. Holmström, Y. Zhang, A. T. Dinkova-Kostova, A. Y. Abramov, Nrf2 regulates ROS production by mitochondria and NADPH oxidase. Biochim Biophys Acta. 1850, 794–801 (2015).

52. A. Pantel, A. Teixeira, E. Haddad, E. G. Wood, R. M. Steinman, M. P. Longhi, Direct type I IFN but not MDA5/TLR3 activation of dendritic cells is required for maturation and metabolic shift to glycolysis after poly IC stimulation. PLoS Biol. 12, e1001759 (2014).

53. D. P. D. Souza, A. Achuthan, M. K. S. Lee, K. J. Binger, M.-C. Lee, S. Davidson, D. L. Tull, M. J. McConville, A. D. Cook, A. J. Murphy, J. A. Hamilton, A. J. Fleetwood, Autocrine IFN-I inhibits isocitrate dehydrogenase in the TCA cycle of LPS-stimulated macrophages. J Clin Invest. 129, 4239–4244 (2019).

54. I. G. Shabalina, M. Yu. Vyssokikh, N. Gibanova, R. I. Csikasz, D. Edgar, A. Hallden-Waldemarson, Z. Rozhdestvenskaya, L. E. Bakeeva, V. B. Vays, A. V. Pustovidko, M. V. Skulachev, B. Cannon, V. P. Skulachev, J. Nedergaard, Improved health-span and lifespan in mtDNA mutator mice treated with the mitochondrially targeted antioxidant SkQ1. Aging (Albany NY). 9, 315–336 (2017).

55. D.-F. Dai, T. Chen, J. Wanagat, M. Laflamme, D. J. Marcinek, M. J. Emond, C. P. Ngo, T. A. Prolla, P. S. Rabinovitch, Age-Dependent Cardiomyopathy in Mitochondrial Mutator Mice is Attenuated by Overexpression of Catalase Targeted to Mitochondria. Aging Cell. 9, 536– 544 (2010).

56. R. H. Hämäläinen, K. J. Ahlqvist, P. Ellonen, M. Lepistö, A. Logan, T. Otonkoski, M. P. Murphy, A. Suomalainen, mtDNA Mutagenesis Disrupts Pluripotent Stem Cell Function by Altering Redox Signaling. Cell Rep. 11, 1614–1624 (2015).

57. J. E. Kolesar, A. Safdar, A. Abadi, L. G. MacNeil, J. D. Crane, M. A. Tarnopolsky, B. A. Kaufman, Defects in mitochondrial DNA replication and oxidative damage in muscle of mtDNA mutator mice. Free Radical Biology and Medicine. 75, 241–251 (2014).

58. A. Saleem, A. Safdar, Y. Kitaoka, X. Ma, O. S. Marquez, M. Akhtar, A. Nazli, R. Suri, J. Turnbull, M. A. Tarnopolsky, Polymerase gamma mutator mice rely on increased glycolytic flux for energy production. Mitochondrion. 21, 19–26 (2015).

59. B. P. Woodall, A. M. Orogo, R. H. Najor, M. Q. Cortez, E. R. Moreno, H. Wang, A. S. Divakaruni, A. N. Murphy, Å. B. Gustafsson, Parkin does not prevent accelerated cardiac aging in mitochondrial DNA mutator mice. JCI Insight. 4 (2019), doi:10.1172/jci.insight.127713.

60. A. Y. Khakoo, M. K. Halushka, J. E. Rame, E. R. Rodriguez, E. K. Kasper, D. P. Judge, Reversible cardiomyopathy caused by administration of interferon alpha. Nat Clin Pract Cardiovasc Med. 2, 53–57 (2005).

61. S. Ioannou, G. Hatzis, I. Vlahadami, M. Voulgarelis, Aplastic anemia associated with interferon alpha 2a in a patient with chronic hepatitis C virus infection: a case report. J Med Case Reports. 4, 268 (2010).

62. J. Chen, Z. Zhang, L. Cai, Diabetic cardiomyopathy and its prevention by nrf2: current status. Diabetes Metab J. 38, 337–345 (2014).

63. J.-M. Lee, K. Chan, Y. W. Kan, J. A. Johnson, Targeted disruption of Nrf2 causes regenerative immune-mediated hemolytic anemia. Proc Natl Acad Sci U S A. 101, 9751– 9756 (2004).

64. K. J. Ahlqvist, R. H. Hämäläinen, S. Yatsuga, M. Uutela, M. Terzioglu, A. Götz, S. Forsström, P. Salven, A. Angers-Loustau, O. H. Kopra, H. Tyynismaa, N.-G. Larsson, K. Wartiovaara, T. Prolla, A. Trifunovic, A. Suomalainen, Somatic Progenitor Cell Vulnerability to Mitochondrial DNA Mutagenesis Underlies Progeroid Phenotypes in Polg Mutator Mice. Cell Metabolism. 15, 100–109 (2012).

65. K. J. Ahlqvist, S. Leoncini, A. Pecorelli, S. B. Wortmann, S. Ahola, S. Forsström, R. Guerranti, C. De Felice, J. Smeitink, L. Ciccoli, R. H. Hämäläinen, A. Suomalainen, MtDNA mutagenesis impairs elimination of mitochondria during erythroid maturation leading to enhanced erythrocyte destruction. Nat Commun. 6, 6494 (2015).

66. J. L. Edmonds, D. J. Kirse, D. Kearns, R. Deutsch, L. Spruijt, R. K. Naviaux, The otolaryngological manifestations of mitochondrial disease and the risk of neurodegeneration with infection. Arch. Otolaryngol. Head Neck Surg. 128, 355–362 (2002).

67. A. Quintana, S. E. Kruse, R. P. Kapur, E. Sanz, R. D. Palmiter, Complex I deficiency due to loss of Ndufs4 in the brain results in progressive encephalopathy resembling Leigh syndrome. PNAS. 107, 10996–11001 (2010).

68. Z. Jin, W. Wei, M. Yang, Y. Du, Y. Wan, Mitochondrial Complex I Activity Suppresses Inflammation and Enhances Bone Resorption by Shifting Macrophage-Osteoclast Polarization. Cell Metabolism. 20, 483–498 (2014).

69. Y. Wang, Q. Liu, T. Liu, Q. Zheng, X. Xu, X. Liu, W. Gao, Z. Li, X. Bai, Early plasma monocyte chemoattractant protein 1 predicts the development of sepsis in trauma patients: A prospective observational study. Medicine (Baltimore). 97, e0356 (2018).

70. H. Matsumoto, H. Ogura, K. Shimizu, M. Ikeda, T. Hirose, H. Matsuura, S. Kang, K. Takahashi, T. Tanaka, T. Shimazu, The clinical importance of a cytokine network in the acute phase of sepsis. Scientific Reports. 8, 13995 (2018).

71. A. J. Lee, B. Chen, M. V. Chew, N. G. Barra, M. M. Shenouda, T. Nham, N. van Rooijen, M. Jordana, K. L. Mossman, R. D. Schreiber, M. Mack, A. A. Ashkar, Inflammatory monocytes require type I interferon receptor signaling to activate NK cells via IL-18 during a mucosal viral infection. J. Exp. Med. 214, 1153–1167 (2017).

72. M. B. Buechler, H. M. Akilesh, J. A. Hamerman, Cutting Edge: Direct Sensing of TLR7 Ligands and Type I IFN by the Common Myeloid Progenitor Promotes mTOR/PI3K-Dependent Emergency Myelopoiesis. The Journal of Immunology (2016), doi:10.4049/jimmunol.1600813.

73. H. M. Akilesh, M. B. Buechler, J. M. Duggan, W. O. Hahn, B. Matta, X. Sun, G. Gessay, E. Whalen, M. Mason, S. R. Presnell, K. B. Elkon, A. Lacy-Hulbert, B. J. Barnes, M. Pepper, J. A. Hamerman, Chronic TLR7 and TLR9 signaling drives anemia via differentiation of specialized hemophagocytes. Science. 363 (2019), doi:10.1126/science.aao5213.

74. G. L. Norddahl, C. J. Pronk, M. Wahlestedt, G. Sten, A. Ugale, M. Sigvardsson, D. Bryder, Accumulating mitochondrial DNA mutations drive premature hematopoietic aging phenotypes distinct from physiological stem cell aging. Cell Stem Cell. 8, 499–510 (2011).

75. R. H. Hämäläinen, J. C. Landoni, K. J. Ahlqvist, S. Goffart, S. Ryytty, M. O. Rahman, V. Brilhante, K. Icay, S. Hautaniemi, L. Wang, M. Laiho, A. Suomalainen, Defects in mtDNA replication challenge nuclear genome stability through nucleotide depletion and provide a unifying mechanism for mouse progerias. Nature Metabolism. 1, 958–965 (2019).

76. K. M. Holmstrom, L. Baird, Y. Zhang, I. Hargreaves, A. Chalasani, J. M. Land, L. Stanyer, M. Yamamoto, A. T. Dinkova-Kostova, A. Y. Abramov, Nrf2 impacts cellular bioenergetics by controlling substrate availability for mitochondrial respiration. Biology Open. 2, 761–770 (2013).

77. R. Erkens, C. M. Kramer, W. Lückstädt, C. Panknin, L. Krause, M. Weidenbach, J. Dirzka, T. Krenz, E. Mergia, T. Suvorava, M. Kelm, M. M. Cortese-Krott, Left ventricular diastolic dysfunction in Nrf2 knock out mice is associated with cardiac hypertrophy, decreased expression of SERCA2a, and preserved endothelial function. Free Radical Biology and Medicine. 89, 906–917 (2015).

78. R. K. Thimmulappa, H. Lee, T. Rangasamy, S. P. Reddy, M. Yamamoto, T. W. Kensler, S. Biswal, Nrf2 is a critical regulator of the innate immune response and survival during experimental sepsis. J. Clin. Invest. 116, 984–995 (2006).

79. A. Cuadrado, A. I. Rojo, G. Wells, J. D. Hayes, S. P. Cousin, W. L. Rumsey, O. C. Attucks, S. Franklin, A.-L. Levonen, T. W. Kensler, A. T. Dinkova-Kostova, Therapeutic targeting of the NRF2 and KEAP1 partnership in chronic diseases. Nat Rev Drug Discov. 18, 295–317 (2019).

80. J. Kwon, E. Han, C.-B. Bui, W. Shin, J. Lee, S. Lee, Y.-B. Choi, A.-H. Lee, K.-H. Lee, C. Park, M. S. Obin, S. K. Park, Y. J. Seo, G. T. Oh, H.-W. Lee, J. Shin, Assurance of mitochondrial integrity and mammalian longevity by the p62–Keap1–Nrf2–Nqo1 cascade. EMBO reports. 13, 150–156 (2012).

81. C. J. Schmidlin, M. B. Dodson, L. Madhavan, D. D. Zhang, Redox regulation by NRF2 in aging and disease. Free Radic. Biol. Med. 134, 702–707 (2019).

82. N. Kubben, W. Zhang, L. Wang, T. C. Voss, J. Yang, J. Qu, G.-H. Liu, T. Misteli, Repression of the Antioxidant NRF2 Pathway in Premature Aging. Cell. 165, 1361–1374 (2016).

83. A. Härtlova, S. F. Erttmann, F. A. Raffi, A. M. Schmalz, U. Resch, S. Anugula, S. Lienenklaus, L. M. Nilsson, A. Kröger, J. A. Nilsson, T. Ek, S. Weiss, N. O. Gekara, DNA damage primes the type I interferon system via the cytosolic DNA sensor STING to promote anti-microbial innate immunity. Immunity. 42, 332–343 (2015).

84. R. Kreienkamp, S. Graziano, N. Coll-Bonfill, G. Bedia-Diaz, E. Cybulla, A. Vindigni, D. Dorsett, N. Kubben, L. F. Z. Batista, S. Gonzalo, A Cell-Intrinsic Interferon-like Response Links Replication Stress to Cellular Aging Caused by Progerin. Cell Reports. 22, 2006–2015 (2018).

85. Q. Yu, Y. V. Katlinskaya, C. J. Carbone, B. Zhao, K. V. Katlinski, H. Zheng, M. Guha, N. Li, Q. Chen, T. Yang, C. J. Lengner, R. A. Greenberg, F. B. Johnson, S. Y. Fuchs, DNA-Damage-Induced Type I Interferon Promotes Senescence and Inhibits Stem Cell Function. Cell Reports. 11, 785–797 (2015).

86. M. F. Manchinu, C. Brancia, C. A. Caria, E. Musu, S. Porcu, M. Simbula, I. Asunis, L. Perseu, M. S. Ristaldi, Deficiency in interferon type 1 receptor improves definitive erythropoiesis in Klf1 null mice. Cell Death & Differentiation. 25, 589–599 (2018).

87. J. Finsterer, M. Frank, Haematological abnormalities in mitochondrial disorders. Singapore Med J. 56, 412–419 (2015).

88. O. Hikmat, C. Tzoulis, C. Klingenberg, M. Rasmussen, C. M. E. Tallaksen, E. Brodtkorb, T. Fiskerstrand, R. McFarland, S. Rahman, L. A. Bindoff, The presence of anaemia negatively influences survival in patients with POLG disease. J. Inherit. Metab. Dis. 40, 861–866 (2017).

